# A New Model of Heart Failure with Preserved Ejection Fraction Induced by Metabolic Syndrome in Ossabaw Miniature Swine

**DOI:** 10.1101/2025.04.29.651284

**Authors:** Xian-Liang Tang, Mouhamad Alloosh, Qinghui Ou, Li Luo, Devendra K. Agrawal, Dinesh K. Kalra, Michael Sturek, Roberto Bolli

**Affiliations:** Institute of Molecular Cardiology, University of Louisville, Louisville, KY 40292; CorVus Biomedical, LLC and CorVus Foundation, Inc; Western University of Health Sciences

**Keywords:** Diastolic dysfunction, hypertension, glucose intolerance, hypercholesterolemia, fibrosis, inflammation, exercise

## Abstract

A major obstacle to progress in heart failure with preserved ejection fraction (HFpEF) is the paucity of clinically relevant animal models. We developed a large, translationally relevant model in Ossabaw minipigs, which are genetically predisposed to the metabolic syndrome (MetS). Pigs were fed a “Western diet” high in calories, fructose, fat, cholesterol, and salt and received 1-2 deoxy-corticosterone acetate (DOCA) depots (n=10). After 6 months, they exhibited liver function abnormalities and marked increases in body weight, arterial blood pressure, serum cholesterol and triglycerides, and plasma glucose and insulin levels (glucose tolerance test), indicating the development of a full MetS. Echocardiography demonstrated no change in LV ejection fraction but progressive concentric LV hypertrophy and left atrial dilatation. Doppler echocardiography showed increased E/e’ ratio and increased peak early (E) and peak late atrial (A) transmitral inflow velocities, with no change in E/A ratio. Right heart catheterization demonstrated increased central venous pressure, pulmonary arterial systolic pressure, and pulmonary capillary wedge pressure. Clinically, pigs exhibited impaired exercise capacity, assessed by treadmill tests, associated with chronotropic incompetence. Pathologic examination showed significant myocardial fibrosis, myocyte hypertrophy, and liver fibrosis. In contrast, lean pigs fed a standard diet (n=3) did not show any changes at 6 months. The Ossabaw porcine model described herein is unique in that it recapitulates the entire constellation of major multiorgan comorbidities and hemodynamic, clinical, and metabolic features of MetS-driven human HFpEF: obesity, arterial hypertension, hyperlipidemia, glucose intolerance, insulin resistance, liver fibrosis and dysfunction, pulmonary hypertension, increased LV filling pressures, concentric LV hypertrophy, LV diastolic dysfunction with preserved systolic function, and impaired exercise capacity. Because of its high clinical relevance, this model is well-suited for exploring the pathophysiology of MetS-driven HFpEF and the efficacy of new therapies.

## INTRODUCTION

Heart failure with preserved ejection fraction (HFpEF) is a common, lethal, disabling, and expensive condition. Its prevalence has reached epidemic proportions (∼3 million in the US and ∼32 million people worldwide; ∼1/2 of all cases of HF), and will continue to rise as obesity and diabetes increase and the population ages [7, 22, 45, 50, 66]. The prognosis of HFpEF is poor, and mortality is at least as high as, if not greater than, that of patients with heart failure with reduced ejection fraction (HFrEF) [7, 22, 45, 50, 66]. Patients with HFpEF have an annual mortality of ∼15%; in those hospitalized with HFpEF, the 5-year mortality is 50-75%, similar to lung cancer [7, 22, 45, 50, 66]. In contrast to HFrEF, there has been little progress in the treatment of HFpEF in the past 40 years [7, 22, 45, 50, 66]. Current therapies (diuretics and SGLT2 inhibitors) have limited efficacy [7, 22, 45, 50, 66]. Thus, HFpEF constitutes a major (and growing) public health problem worldwide, a leading cause of morbidity and mortality, and an increasing burden on healthcare systems; it will soon become one of the greatest challenges in healthcare [7, 22, 45, 50, 66]. In 2017, the NHLBI Working Group on HFpEF concluded that the treatment of HFpEF is “the major unmet need in cardiovascular medicine today” [50].

Among the various subtypes of syndromes within the spectrum of human HFpEF (e.g., HFpEF induced by hypertension, renal disease, aging), the most prevalent is HFpEF driven by the metabolic syndrome (MetS), which accounts for 60-70% of all cases and is associated with obesity and hyperglycemia/insulin resistance [7, 10, 22, 45, 50]. A major obstacle to progress in both the pathophysiology and therapy of MetS-induced HFpEF has been the lack of translationally relevant animal models, due to the difficulty in recapitulating the systemic, multiorgan dysfunction and comorbid-laden phenotype of patients with this syndrome [10, 50, 68]. Indeed, the aforementioned Working Group identified development of improved large animal models of HFpEF that incorporate comorbid conditions as a “research priority” [50]. Most existing models of MetS-induced HFpEF use rodents, which are less relevant to the human syndrome and not suitable for testing device-based therapies.

The objective of this study was to develop a large animal model of MetS-induced HFpEF that recapitulates the key elements of the clinical syndrome, including impaired exercise capacity, and can be used for preclinical studies of new therapies. To this end, we used Ossabaw miniature pigs, which are genetically predisposed to MetS, mimicking the constellations of comorbidities and multiorgan abnormalities associated with human MetS [10, 12, 55, 68]. The observations reported herein describe a new animal model of HFpEF induced by diet and deoxy-corticosterone acetate (DOCA) that exhibits both hemodynamic (increased left ventricular [LV] filling pressures) and clinical (exercise intolerance) manifestations of diastolic dysfunction and incorporates the major comorbidities and the systemic, multiorgan dysfunction associated with human HFpEF, and thus has high clinical relevance. This is the only porcine model of HFpEF published heretofore that exhibits all of the following components: a full MetS, a full metabolic characterization, liver fibrosis and dysfunction, and reduced exercise tolerance.

## METHODS

All animal experiments were performed in accordance with *the Guide for the Care and Use of Laboratory Animals* published by the US National Institutes of Health (Eighth Edition, Revised 2010) and with the guidelines of the Animal Care and Use Committees of the University of Louisville, School of Medicine and CorVus Biomedical, LLC. Ossabaw miniature swine of either sex (age, 22.1 ± 1.8 months; body weight, 48.6 ± 1.1 kg) were used. At this age, Ossabaw pigs have reached sexual maturity and their body weight is nearly a plateau [52, 55]. The relatively steady body weight facilitates repeated physiological and metabolic assessments in lean pigs to compare with pigs on a “Western diet” [55]. Throughout the studies, pigs were housed in the Ossabaw pig facility at CorVus Biomedical, LLC, in Crawfordsville, IN.

### Development of HFpEF

Pigs were assigned to a lean control cohort or a HFpEF cohort **(Fig. 1)**. In the lean cohort, pigs were fed a standard diet (2,300 kcal/day) given once a day (LabDiet Mini-Pig Grower 5L80 [LabDiet.com]) composed of (in %kcal): 18% protein, 11% fat, 71% carbohydrates, 0% cholesterol, and 3.75 g of salt. The lean healthy pigs were used as a reference control group to assess the development of abnormalities in the HFpEF cohort during the 6-month study.

**Fig. 1.**
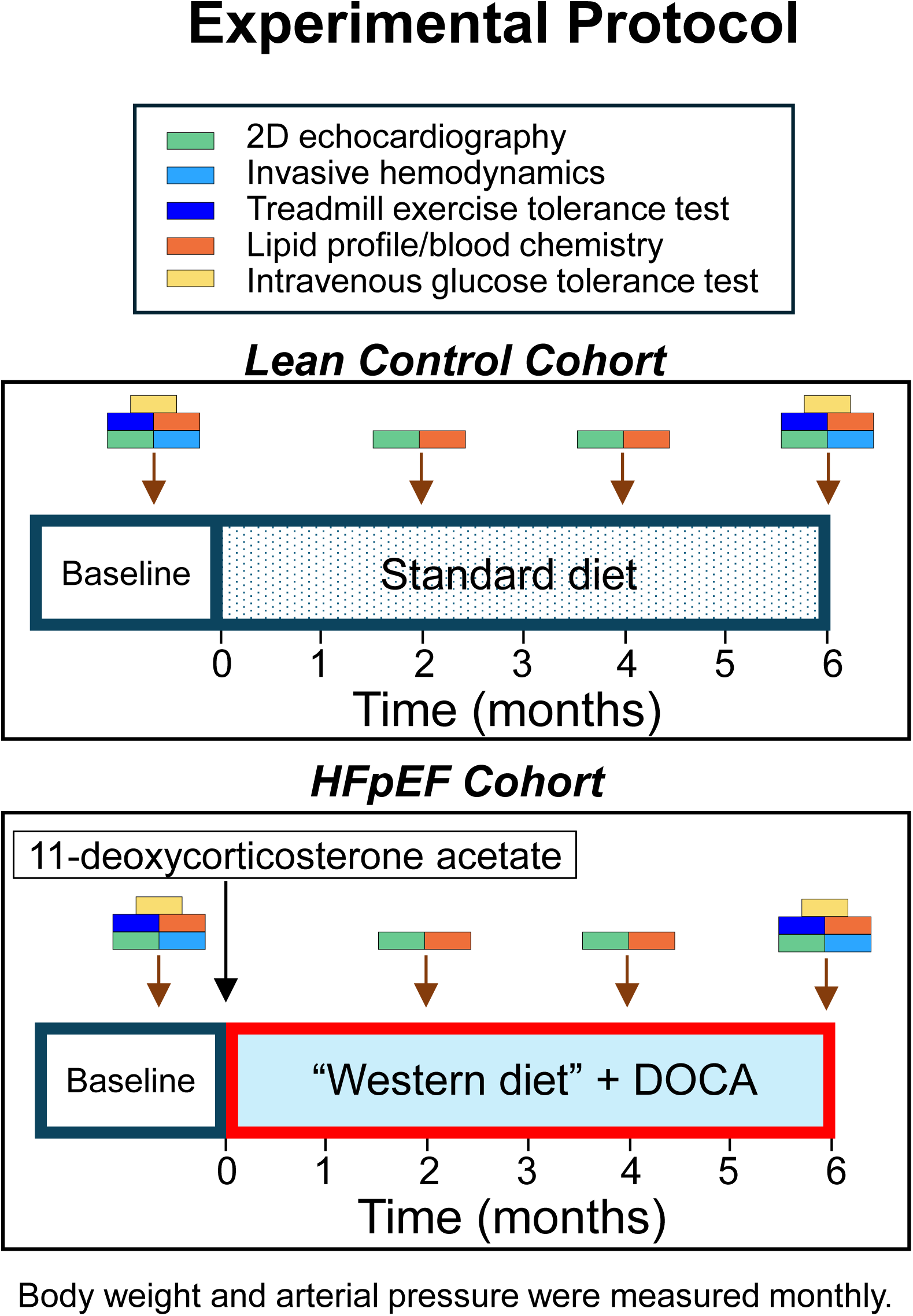
Experimental protocol.

In the HFpEF cohort, pigs were fed a “Western diet” high in calories (4,600 kcal/day, 1 kg/day), fructose, fat, cholesterol, and salt. Specifically, this diet consisted of (in % kcal): 16.3% protein, 40.8% carbohydrates (19% kcal fructose), and 42.9% fat. Fat calories were derived from a mixture of lard, hydrogenated soybean oil, and hydrogenated coconut oil. The diet was supplemented with 2.0% cholesterol by weight and 20 g of salt. This diet is a variation of the diet specially formulated for Ossabaw miniature swine and mixed by CorVus Biomedical, LLC (KT324, Purina Test Diet, Richmond, IN). This diet is well established to induce MetS characterized by obesity, insulin resistance, glucose intolerance, dyslipidemia, and hypertension in Ossabaw pigs [2, 10, 12, 35, 55, 68]. In addition, pigs in the HFpEF cohort received a subcutaneous DOCA depot (50 mg/kg, 200 mg pellets, 60-day release) at baseline; in 5 pigs, the DOCA depot was repeated 3 months later to ensure a robust hypertensive phenotype [49, 52]. The DOCA pellets were implanted while the pig was anesthetized for echocardiographic studies. Pigs were maintained on the Western diet for the 6-month period described in this report.

In both cohorts, an extensive battery of physiologic and metabolic analyses was performed **(Fig. 1)**. After 6 months, pigs were kept on a modified Western diet without DOCA for 5 months and enrolled in pilot studies of experimental therapies (not reported here due to small group sizes). All surviving pigs were euthanized 11 months after the start of the study and pathologic analyses were performed. Only control animals that did not receive any therapy are included in the pathologic results reported here.

### Noninvasive blood pressure measurements

Using a low-stress restraint sling [42], noninvasive measurements of blood pressure were obtained in conscious swine every month with a sphygmomanometer on a limb or tail [12, 13, 56]. Pigs were first acclimated to the sling for 4-6 sessions lasting 10-30 minutes on separate days to ensure that blood pressures were not due to the initial stress of the restraint. Blood pressure was measured every 2-5 minutes for at least 4 measurements over a 20-min period, and data were averaged.

### Intravenous glucose tolerance tests (IVGTTs)

Conscious swine were fasted overnight and placed in a low-stress restraint sling [42]. A 50% glucose solution was given intravenously (1 g glucose/kg body weight) via a percutaneously placed jugular venous catheter. Blood samples (3 mL) were taken at time 0 (just before glucose injection) and at 5, 10, 20, 30, 40, 50, and 60 min thereafter [12, 39, 42, 56]. Blood glucose values were monitored by use of a ReliOn Premier glucose meter. Plasma insulin was quantified using an ELISA kit (Sigma-Aldrich St. Louis, MO) at the Indiana University School of Medicine Diabetes Research Core.

### Lipid profile and blood chemistry

Serum samples were assayed for total, LDL, and HDL cholesterol, triglycerides, blood urea nitrogen, creatinine, albumin, globulin, electrolytes, creatine phosphokinase, ALT, AST, GGT, and alkaline phosphatase (Antech Diagnostics, West Lafayette, IN).

### Echocardiography

Echocardiographic studies were performed as described [1, 62] at baseline and 2, 4, and 6 months later. Briefly, pigs were lightly anesthetized (isoflurane, 0.5-1.5%) and rested on a pig sling in the supine position or tilted 10-20 degrees towards the right lateral position in order to obtain optimal images. Using a portable Chison SonoBook 9 ultrasound machine and a P2-V or a P5-V phase array probe, transthoracic 2-D (B-mode) parasternal long axis images and M-mode images were acquired [24, 26, 63]. LV volumes, LVEF, left atrial area, and left atrial fractional area change were measured using the 2-D long axis images [12, 52]. LV wall thickness and cavity diameter were measured from the M-mode view at the LV mid-papillary muscle level [12, 52]. Pulse-wave Doppler images were obtained at the mitral valve orifice with a maximal corrected angle of 40 degrees for assessment of transmitral inflow velocities [12, 52]. Pulse-wave tissue Doppler images were obtained at both the medial and lateral annulus of the mitral valve with a maximal corrected angle of 40 degrees for calculation of tissue velocities during early LV filling (é) at the mitral valve annulus [12, 52]. All images were acquired and analyzed by a single researcher (XLT) and all data were analyzed using the Chison SonoBook 9 onboard imaging software.

### Hemodynamic studies

Direct measurement of cardiac filling pressures is the gold standard for the diagnosis of HFpEF [7, 22, 45, 50]. Hemodynamic studies were performed at baseline and 6 months. Briefly, under isoflurane anesthesia (1-2.5%), a 9F sheath was inserted percutaneously into the right external jugular vein and a Swan-Ganz catheter (Edwards Lifesciences) was introduced via the 9F sheath into the superior vena cava and the right atrium. After the balloon was inflated, the catheter was floated to the right ventricle and the pulmonary artery, and then further advanced into a pulmonary artery branch until pressures dropped suddenly, indicating that the balloon at the catheter tip was wedged **(Fig. 6)**. Central venous pressure, pulmonary artery systolic, diastolic, and mean pressure, and pulmonary capillary wedge pressure were measured (Meritrans DTX Pressure Transducer System, Merit Medical). Pressure signals were recorded and measured using LabChart 8 Software, ADInstruments.

### Treadmill tests

Impaired exercise tolerance is the clinical hallmark of HFpEF [7, 22, 45, 50, 68]. Submaximal exercise stress tests were performed using protocols similar to those used in previous studies **(Fig. 7)** [4, 6, 13, 44, 54]. Pigs were acclimated to the treadmill [44] by slow walking (3.5 km/h, 5-10 min) for 3-4 sessions. After suitable acclimation sessions (typically 3-4), the pigs were fitted with a Polar H10 heart rate (HR) monitor placed around the thorax for the stress test. Resting HR was first recorded standing on the treadmill. The exercise consisted of a warm-up walk at 3.5 km/h and 0% grade for 1 min and at 6.1 km/h for 1 min, followed by a 10-min fast walk at 7.7 km/h, which elicits a HR of ∼120-160/min (∼50-65% of maximum) in healthy, lean pigs. The treadmill speed was then decreased to 3.5 km/h for 2 min to observe recovery of HR and then stopped for 2 min to determine whether there was further recovery toward resting, pre-exercise HR **(Fig. 7)**. The submaximal HR during the 10-min fast walk was recorded at baseline to establish normal exercise tolerance and reliability of the test. The test was repeated after 6 months. Increased exercise HR at 6 months compared to baseline would indicate decreased exercise capacity [4, 6, 13, 44, 54]. Inability to complete the 10-min test (abnormal gait, labored breathing, excessive vocalization, excessive HR) would indicate decreased exercise tolerance. In addition, decreased or unchanged exercise HR compared to baseline and the lean control group would indicate chronotropic incompetence, which has been noted in HFpEF [15, 21, 34, 43, 44, 72]. We used the mean HR during minutes 4-8 of the 7.7 km/h fast walk stage of the treadmill test to obtain steady-state HR responses to the workload.

### Histologic studies

At the conclusion of the study, pigs were euthanized and LV samples harvested for pathologic analysis. The protocols for histologic analyses have been described [57–62, 64]. Briefly, frozen LV tissue samples were sectioned in 8-µm thick sections and mounted on glass slides. Liver samples were processed in a similar fashion. Histologic and immunofluorescent stainings were performed following manufacturer’s protocols. All images were captured using a Nikon Eclipse 80i fluorescence microscope (Nikon, Tokyo, Japan) with a ×4 objective. Representative images were taken with the same ×4 objective, unless otherwise specified. Digital imaging and quantifications were performed using NIS Elements Software (Nikon, Tokyo, Japan). Myocardial and liver collagen content was quantitated using Masson’s trichrome staining. Tissue fibrosis was quantified by calculating the percentage of total Masson’s trichrome positive tissue (blue) over the total tissue area using Image J software (National Institutes of Health, Bethesda, Maryland). Rhodamine-conjugated wheat germ agglutinin (WGA, Vector Labs, Newark, CA) staining was performed to outline cell membranes and facilitate measurements of cardiomyocyte cross-sectional area. Myocytes were stained with α-sarcomeric actin antibody (α-SA) and nuclei were stained with DAPI (4’,6-diamidino-2-phenylindole). Representative images were taken with the ×20 objective and NIS Elements Software was used to measure the cross-sectional area of 140-150 cells per animal.

### Statistical analysis

The major endpoints of this study were cardiac filling pressures, which were assessed by their clinical manifestation (exercise tolerance) and by hemodynamic (pulmonary capillary wedge pressure) and echocardiographic (E/é ratio) parameters [45, 50]. Body weight, arterial blood pressure, serum lipid levels, and echocardiographic parameters were compared using a two-way (time and group) repeated-measures ANOVA, followed by *post hoc* contrasts (Tukey’s *t*-tests or Student’s *t*-tests with Bonferroni correction) [40, 48]. IVGTT results (serum glucose and insulin levels) were compared within the HFpEF group with a two-way (test and time) repeated-measures ANOVA. For the other variables, Student’s t-tests (unpaired or paired as appropriate) with Bonferroni correction were used to evaluate diet effects at the same time point, as well as to evaluate time effect within the same diet group. All analyses were conducted with SigmaStat 3.5 for Windows (Systat Software Inc. Richmond, California). Data are expressed as means ± SEM. A *P* value <0.05 was considered significant.

## RESULTS

### Development of metabolic syndrome

Ten pigs (5 males, 5 females) were assigned to the Western diet + DOCA (HFpEF group) and 3 (2 males, 1 female) to the regular diet (lean control group) **(Fig. 1)**. For various logistical and technical reasons, not all tests could be performed in all pigs. In lean controls (n=3), measurements of blood pressure, serum lipids, and right heart hemodynamics were performed in 2 pigs and IVGTTs in 1 pig. In the HFpEF cohort (n=10), IVGTTs were performed in 8 pigs and hemodynamic studies and treadmill exercise tests in 9 pigs. Histologic evaluation was performed in 2 lean controls and 6 HFpEF pigs.

As shown in **Fig. 2**, body weight remained relatively constant in lean controls over 6 months. In contrast, pigs on a Western diet + DOCA exhibited a progressive, marked increase in body weight that tended to plateau at 6 months. Arterial blood pressure remained within the normal range in lean controls, but increased progressively in the HFpEF group, again plateauing at ∼5 months **(Fig. 2)**. The development of obesity and hypertension was associated with a marked increase in serum cholesterol and triglycerides levels, which reached a plateau at 2 months and remained relatively constant over the next 4 months **(Fig. 3**; **Table 1)**. The increase in total cholesterol was largely accounted for by an increase in LDL cholesterol (LDL-C), although HDL cholesterol also increased **(Fig. 3)**. IVGTTs were performed at baseline and 6 months later **(Fig. 4)**. In lean controls, the response to the glucose load was similar at baseline and 6 months later **(Figs. 4A and C)**. In contrast, in the HFpEF group, following the intravenous bolus of glucose plasma levels of both glucose and insulin were higher at 6 months compared with baseline **(Figs. 4B and D)**, indicating impaired glucose tolerance and insulin resistance.

**Fig. 2.**
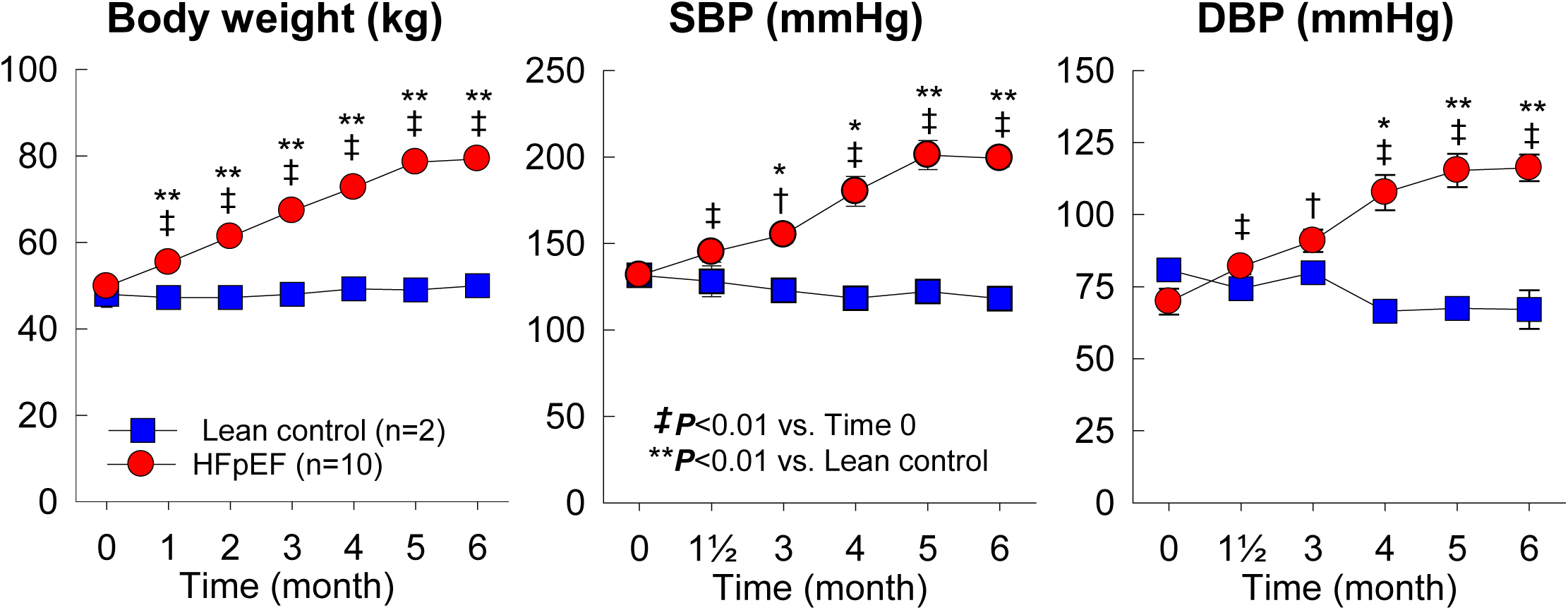
Body weight, systolic blood pressure (SBP), and diastolic blood pressure (DBP) in HFpEF pigs and lean controls. By 6 months, body weight Increased by 58%, SBP by 51%, and DBP by 67% vs. time 0 (baseline) in HFpEF pigs but remained stable in lean controls. Data are mean ± SEM.

**Fig. 3.**
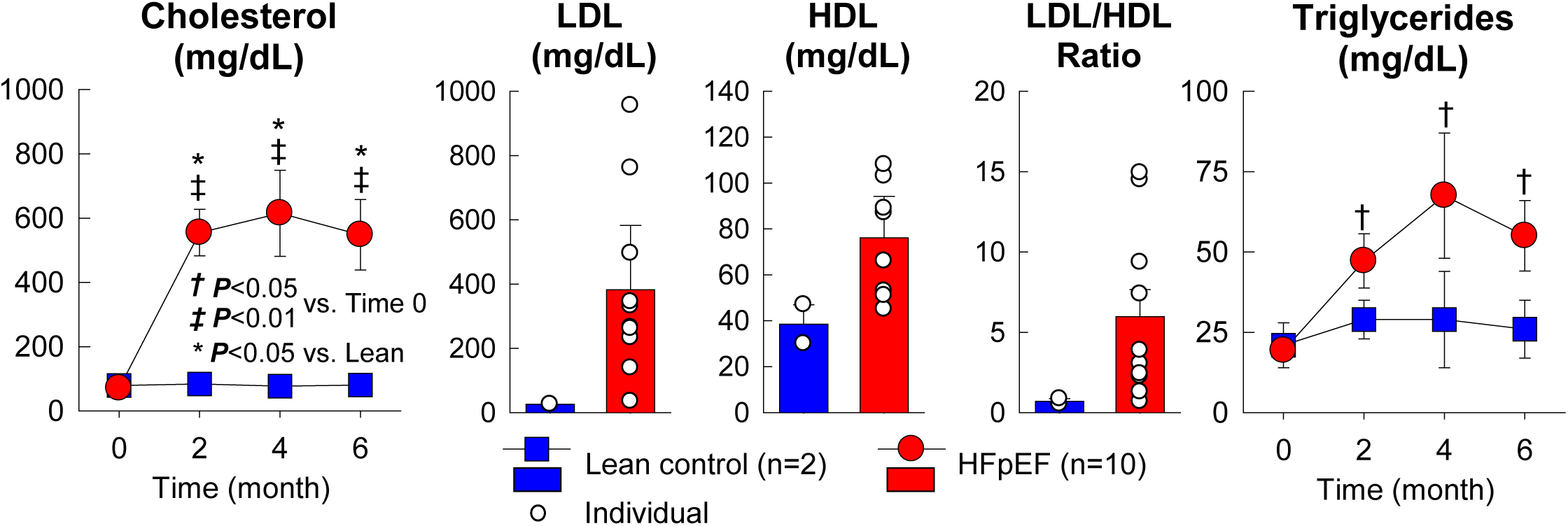
Serum levels of cholesterol and triglycerides in HFpEF pigs and lean controls. In HFpEF pigs, both variables increased at 2 months and remained elevated thereafter. LDL, low-density cholesterol; HDL, high-density cholesterol. Data are mean ± SEM; open circles represent individual pigs.

**Fig. 4.**
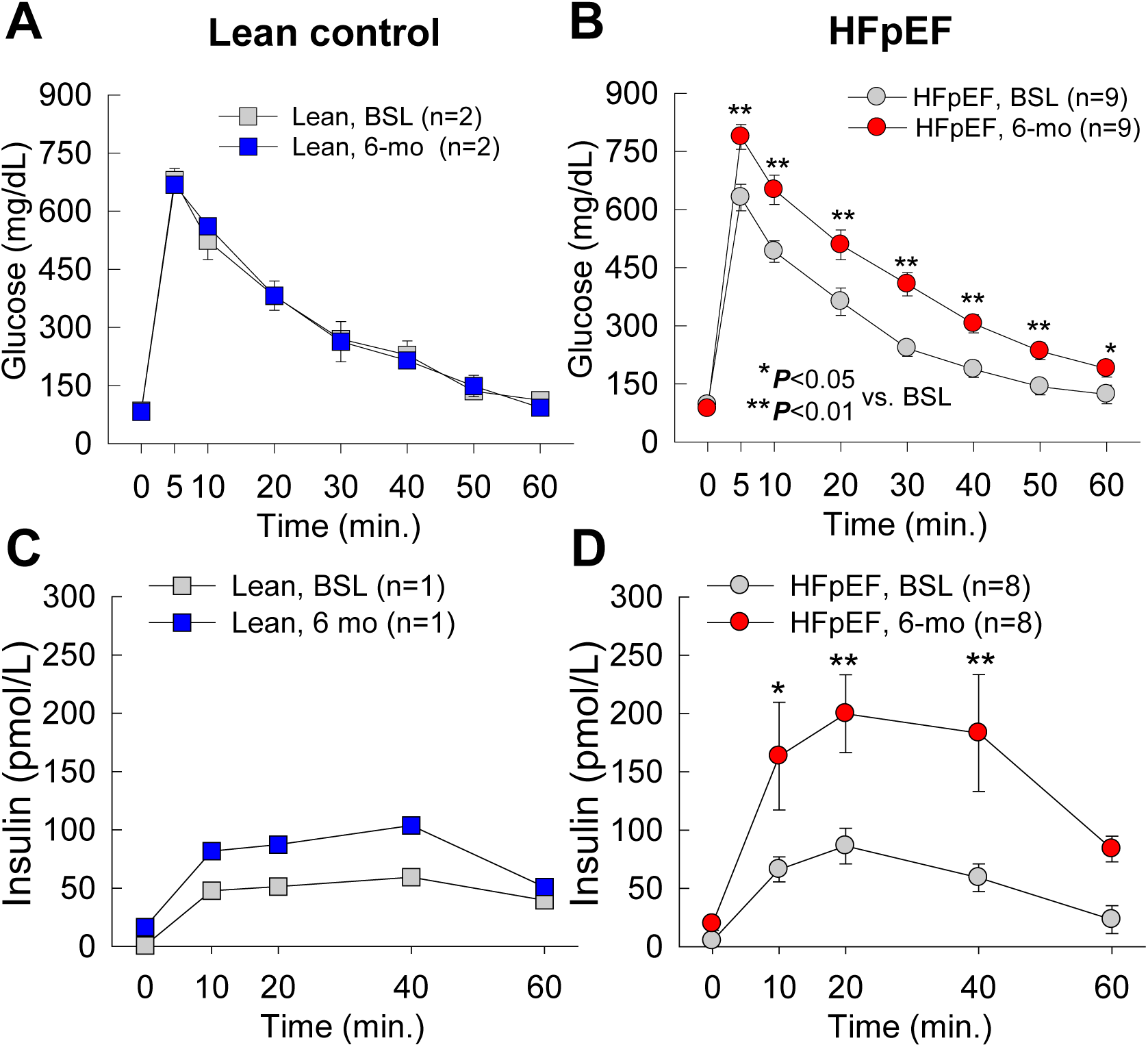
Intravenous glucose tolerance tests at baseline (BSL) and after 6 months of Western diet plus DOCA (HFpEF pigs) or regular diet (lean control). In HFpEF pigs, plasma glucose levels **(<u>top panels</u>)** and insulin levels **(<u>bottom panels</u>)** were significantly higher at 6 months vs. baseline, indicating insulin resistance and impaired glucose tolerance. Horizontal axis indicates time (min) after injection of glucose bolus. Data are mean ± SEM.

**Table 1.** Blood chemistry.

|  | Time | Lean control (n=2) |  | HFpEF (n=9) |  |
| --- | --- | --- | --- | --- | --- |
|  |  | Mean | SEM | Mean | SEM |
| **Total Protein**<br>**(g/dL)** | Baseline | 7.03 | 0.33 | 7.30 | 0.18 |
|  | 2-month | 6.77 | 0.39 | 6.16 | 0.12<sup>b</sup> |
|  | 4-month | 6.75 | 0.15 | 6.82 | 0.11<sup>a</sup> |
|  | 6-month | 6.75 | 0.45 | 6.74 | 0.14<sup>b</sup> |
| **Albumin**<br>**(g/dL)** | Baseline | 3.97 | 0.09 | 4.14 | 0.09 |
|  | 2-month | 3.57 | 0.13 | 3.53 | 0.06<sup>b</sup> |
|  | 4-month | 3.55 | 0.25 | 3.64 | 0.06<sup>b</sup> |
|  | 6-month | 3.60 | 0.60 | 3.43 | 0.14<sup>b</sup> |
| **Globulin**<br>**(g/dL)** | Baseline | 3.07 | 0.27 | 3.16 | 0.17 |
|  | 2-month | 3.20 | 0.26 | 2.63 | 0.09 |
|  | 4-month | 3.20 | 0.10 | 3.18 | 0.10 |
|  | 6-month | 3.15 | 0.15 | 3.31 | 0.18 |
| **AST (SGOT,**<br>**IU/L)** | Baseline | 40.00 | 8.08 | 30.00 | 1.64 |
|  | 2-month | 28.33 | 5.36 | 64.60 | 5.89<sup>b**</sup> |
|  | 4-month | 33.00 | 12.00 | 75.70 | 15.83<sup>a</sup> |
|  | 6-month | 82.00 | 53.00 | 50.90 | 5.98<sup>b</sup> |
| **ALT**<br>**(SGPT, IU/L)** | Baseline | 35.00 | 7.23 | 33.20 | 3.08 |
|  | 2-month | 37.33 | 6.39 | 39.60 | 5.04 |
|  | 4-month | 36.00 | 1.00 | 42.40 | 5.13 |
|  | 6-month | 44.00 | 6.00 | 34.40 | 5.91 |
| **Alk**<br>**Phosphatase**<br>**(IU/L)** | Baseline | 49.33 | 17.57 | 62.80 | 7.39 |
|  | 2-month | 51.00 | 11.79 | 138.00 | 32.28<sup>a</sup> |
|  | 4-month | 45.00 | 1.00 | 207.00 | 63.78<sup>a</sup> |
|  | 6-month | 46.00 | 4.00 | 141.10 | 32.37<sup>a</sup> |
| **GGT**<br>**(IU/L)** | Baseline | 42.67 | 3.53 | 47.70 | 6.20 |
|  | 2-month | 44.67 | 7.54 | 53.20 | 5.73 |
|  | 4-month | 43.00 | 2.00 | 75.20 | 6.66<sup>b</sup> |
|  | 6-month | 49.00 | 6.00 | 86.50 | 8.59<sup>b</sup> |
| **BUN**<br>**(mg/dL)** | Baseline | 15.67 | 2.33 | 14.10 | 0.43 |
|  | 2-month | 16.33 | 0.33 | 12.80 | 1.54 |
|  | 4-month | 17.00 | 2.00 | 12.60 | 0.87 |
|  | 6-month | 15.00 | 2.00 | 11.90 | 0.71 |
| **Creatinine**<br>**(mg/dL)** | Baseline | 1.07 | 0.09 | 1.10 | 0.05 |
|  | 2-month | 1.13 | 0.09 | 0.83 | 0.06<sup>a</sup> |
|  | 4-month | 1.25 | 0.25 | 0.89 | 0.07<sup>a</sup> |
|  | 6-month | 1.10 | 0.10 | 1.00 | 0.04 |
| **Cholesterol**<br>**(mg/dL)** | Baseline | 74.00 | 6.81 | 70.40 | 5.06 |
|  | 2-month | 75.67 | 8.41 | 554.50 | 72.23<sup>b*</sup> |
|  | 4-month | 77.00 | 1.00 | 614.70 | 133.96<sup>b*</sup> |
|  | 6-month | 80.50 | 11.50 | 547.80 | 109.66<sup>b*</sup> |
| **Triglyceride**<br>**(mg/dL)** | Baseline | 18.00 | 5.03 | 19.30 | 2.24 |
|  | 2-month | 24.00 | 6.08 | 47.20 | 8.40<sup>b</sup> |
|  | 4-month | 69.00 | 55.00 | 67.50 | 19.49<sup>a</sup> |
|  | 6-month | 26.00 | 9.00 | 55.00 | 10.94<sup>b</sup> |
| **Amylase**<br>**(IU/L)** | Baseline | 854.67 | 164.90 | 1278.20 | 110.71 |
|  | 2-month | 1060.00 | 367.28 | 951.10 | 121.13<sup>b*</sup> |
|  | 4-month | 1056.50 | 140.50 | 978.50 | 128.26<sup>b*</sup> |
|  | 6-month | 1311.00 | 476.00 | 969.90 | 147.23<sup>b*</sup> |

<sup>a</sup> $P < 0.05$ , <sup>b</sup> $P < 0.01$ vs. baseline within group; \* $P < 0.05$ , \*\* $P < 0.01$ vs. lean control

Liver function tests showed increases in AST, GGT [69], and alkaline phosphatase in the HFpEF cohort but not in the lean cohort **(Table 1)**, suggesting the development of metabolic dysfunction-associated steatotic liver disease (MASLD) [8]. The decrease in albumin observed in HFpEF pigs **(Table 1)** is also consistent with this concept. Amylase levels decreased significantly in HFpEF pigs **(Table 1)**, which is consistent with the presence of MetS and may suggest chronic pancreatitis [38].

### Echocardiographic studies

Serial echocardiographic studies were conducted at baseline and then at 2, 4, and 6 months **(Fig. 1)**; the results are summarized in **Fig. 5** and **Supplementary Table 1.** In lean controls, all parameters remained relatively constant throughout the study. In contrast, in the HFpEF cohort there was a progressive increase in LV wall thickness, both in the interventricular septum and in the posterior wall, associated with a parallel increase in LV mass and in relative wall thickness, indicating the development of concentric LV hypertrophy. Left atrial area also increased progressively while left atrial area change during diastole decreased, suggesting elevated LV filling pressures. LV ejection fraction (EF), calculated by the Simpson’s method, remained normal throughout the 6-months period. Doppler flow echocardiography showed an increase in both early (E) and late atrial (A) transmitral inflow velocities; as a result, the E/A ratio did not change significantly during the 6-month interval **(Fig. 5)**. The ratio of E to medial and lateral early diastolic mitral annular tissue velocity e’ (E/e’) is used clinically to diagnose HFpEF [7, 22, 45, 68]. Both medial and lateral E/e’ ratios increased; the average E/e’ ratio was significantly elevated at 2 months and continued to increase thereafter **(Fig. 5, Supplementary Table 1)**, indicating rapid development of diastolic dysfunction and HFpEF.

**Fig. 5.**
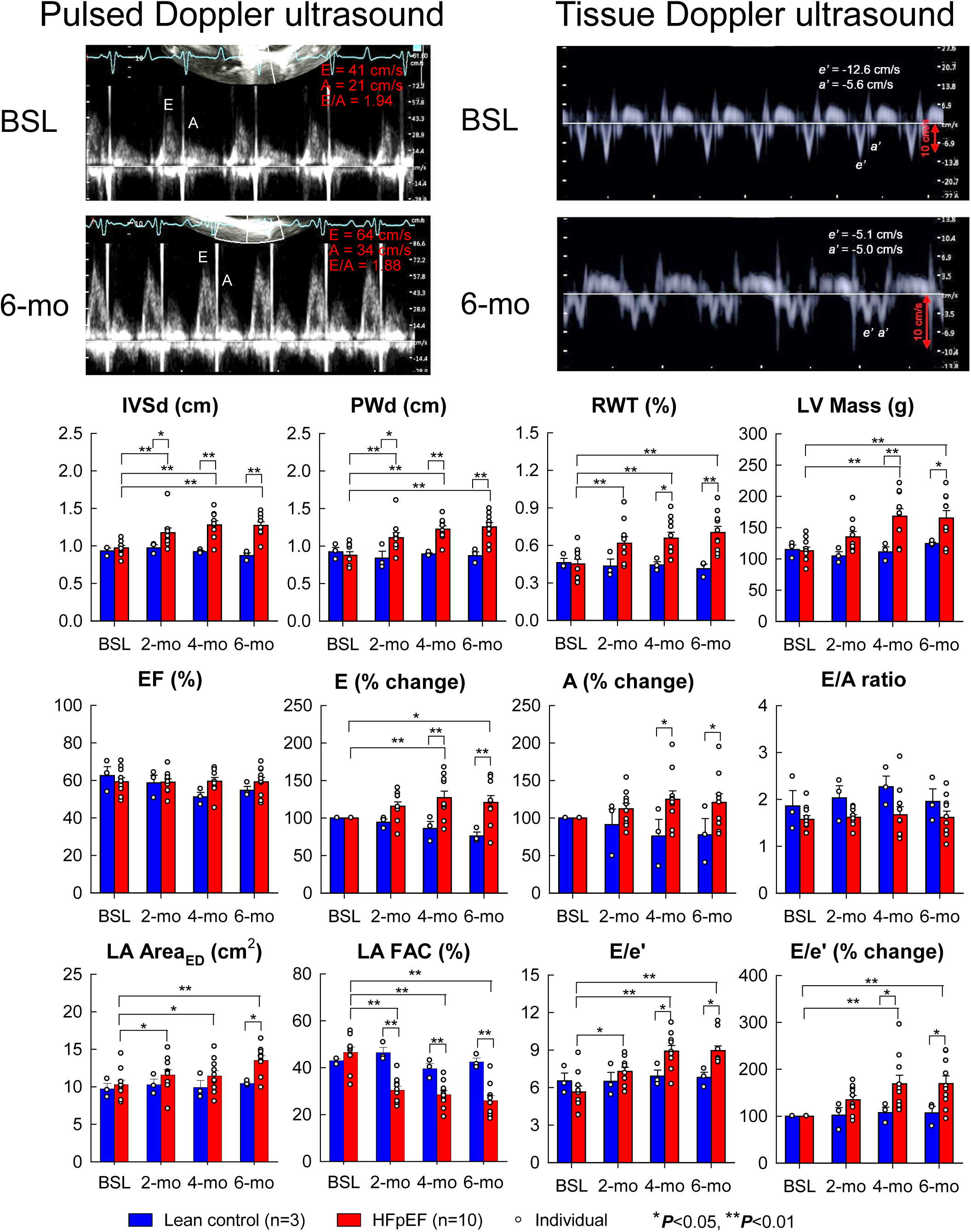
Echocardiographic data in HFpEF and lean control pigs at time 0 (baseline [BSL]) and 2, 4, and 6 months later (2-mo, 4-mo, 6-mo). The images at the top show examples of pulsed Doppler ultrasound and tissue Dopler ultrasound tracings at the mitral valve level at baseline and 6 months later in a HFpEF pig. Note the different scale for tissue Doppler tracings at baseline vs. 6 months (red arrows); in this pig, e’ velocity decreased from -12.6 cm/s to -5.1 cm/s. **<u>Top panels</u>:** Interventricular septum thickness (IVSd), posterior LV wall thickness (PWd), relative LV wall thickness (RWT), and LV mass. (Relative wall thickness [RWT] is calculated as twice the posterior wall thickness divided by the LV end-diastolic diameter; a normal RWT range is generally considered to be between 0.32 and 0.42.) **<u>Middle panels:</u>** LV ejection fraction (EF), peak E velocity (% change from baseline), peak A velocity (% change from baseline), and E/A ratio. **<u>Bottom panels</u>:** Left atrial area (LA area), LA fractional area change (LA FAC), average medial and lateral E/e’ ratio (absolute value), and average medial and lateral E/e’ ratio (% change from baseline). All variables were stable in lean controls. In contrast, HFpEF pigs exhibited progressive LV hypertrophy and left atrial dilatation and a progressive increase in E, A, and E/e’ with no change in EF. These changes were already evident at 2 months of Western diet and became more prominent with time. Data are mean ± SEM; open circles represent individual pigs.

### Hemodynamic studies

At baseline, right heart catheterization showed pressures within normal limits in both groups **(Fig. 6, Supplementary Table 2)**. After 6 months, however, in the HFpEF cohort central venous pressure was elevated and pulmonary artery systolic pressure and pulmonary capillary wedge pressure were significantly increased compared with baseline values. These changes did not occur in lean controls **(Fig. 6)**.

**Fig. 6.**
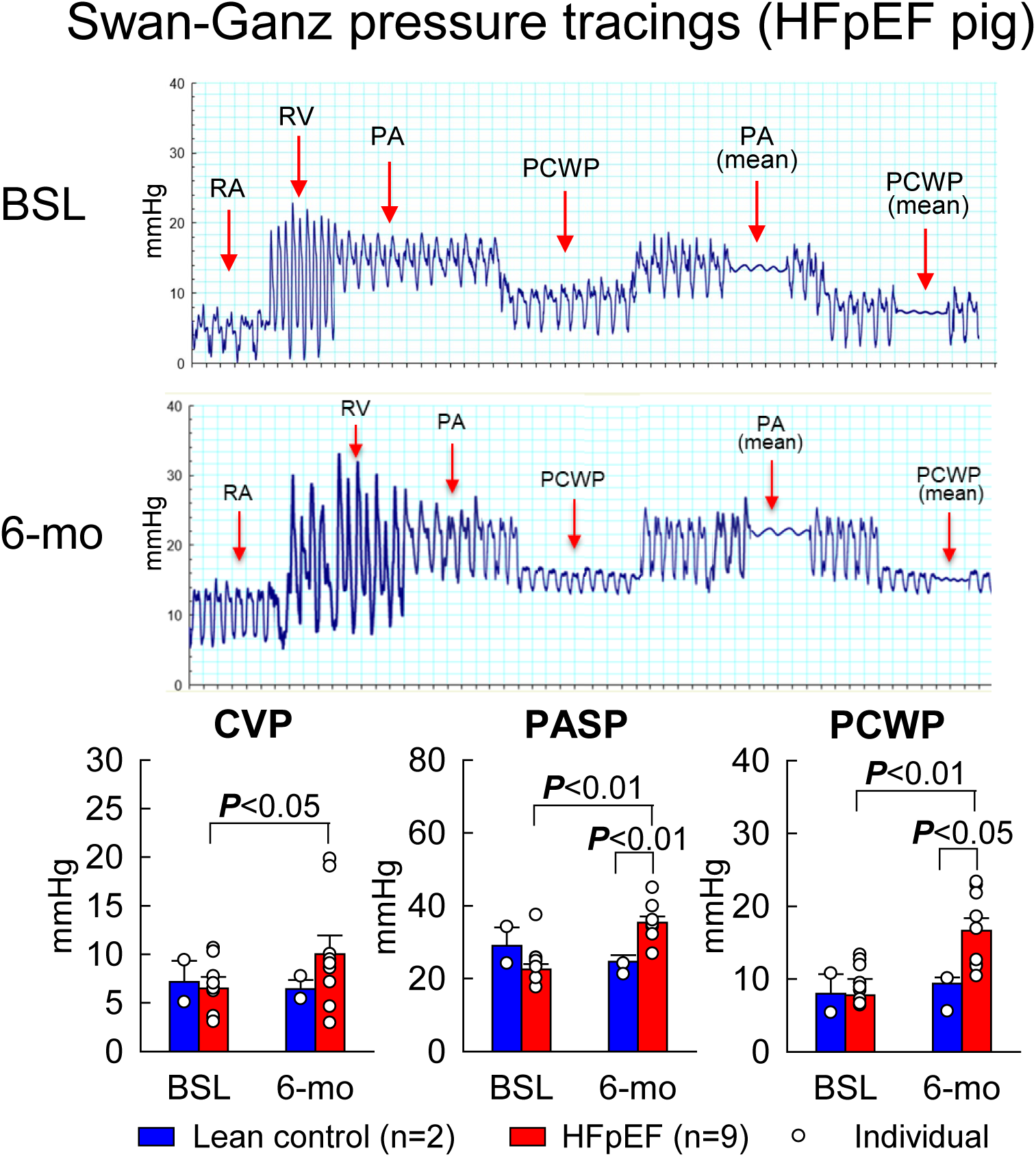
Hemodynamic parameters (central venous pressure [CVP], pulmonary artery systolic pressure [PASP], and pulmonary capillary wedge pressure [PCWP]) in HFpEF and lean control pigs at baseline and 6 months later. In HFpEF pigs, all three hemodynamic variables were increased at 6 months vs. baseline. The top panels illustrate representative pressure recordings obtained with a Swan-Ganz catheter in a HFpEF pig at baseline and 6 months later. RA, right atrium; RV, right ventricle; PA, pulmonary artery. Data are mean ± SEM; open circles represent individual pigs.

### Exercise tolerance

When HFpEF pigs were subjected to treadmill exercise, the duration of exercise tolerated by the animals was decreased significantly, by 37% at 6 months compared with baseline (from 10.0±0 min to 6.3±0.9 min; *P*<0.005) **(Fig. 7)**. Only 2 of 9 HFpEF pigs were able to complete the 10-min fast walking stage of the test at 6 months, indicating impaired exercise tolerance. In 2 pigs the test was discontinued because of dangerously high HR of 199 and 179 bpm, which were 2 standard deviations above the mean HR for the lean, healthy pigs at baseline. In the remaining 7 pigs, average exercise HR during the fast-walking stage did not increase vs. baseline. This is contrary to the typical increase in exercise HR that would be expected given the increased metabolic demand of obesity. Instead, in these 7 HFpEF pigs the average exercise HR decreased from 131±8 b/min at baseline to 117±3 b/min at 6 months (*P*<0.05) and was numerically lower than in lean animals for the entire duration of the 10-min fast walking stage **(Fig. 7)**, indicating chronotropic incompetence. In contrast, after 6 months there was no change in either exercise duration or HR in lean controls compared with baseline **(Fig. 7)**.

**Fig. 7.**
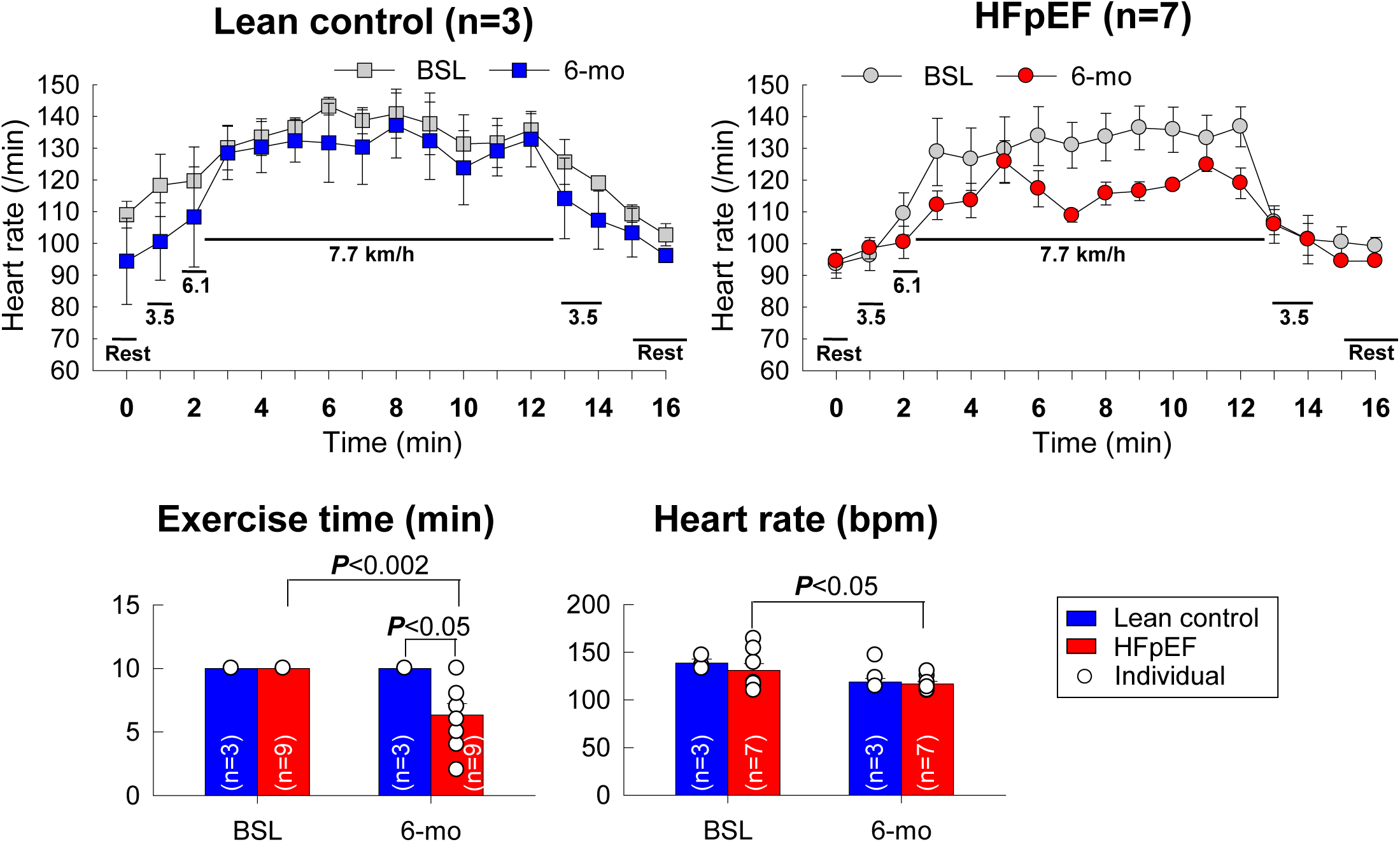
Exercise duration and HR during exercise in HFpEF and lean control pigs at baseline (BSL) and 6 months later. The top panels illustrate HR at the various stages of the exercise protocol at baseline and 6 months later. Note that throughout the 10-min fast walking stage (the main stage of the exercise test), HR was lower at 6 months vs. baseline in HFpEF pigs but not in lean pigs; during this stage, the average HR was significantly lower than at baseline in HFpEF swine **(bottom right panel)**. In addition, exercise duration decreased significantly (-37%) at 6 months vs. baseline in HFpEF pigs but not in lean controls **(bottom left panel)**. Data are mean ± SEM; open circles represent individual pigs.

### Pathologic analysis

At the end of the study (after 11 months of normal or Western diet), Masson’s trichrome-stained LV sections were analyzed to determine the effect of the MetS on myocardial collagen deposition. As shown in **Figs. 8A and B**, myocardial collagen content was markedly increased in HFpEF pigs vs. lean controls (4.6±0.7 % of LV area vs. 1.3±0.5 %, *P*<0.05), indicating significant myocardial fibrosis. WGA staining enabled us to accurately measure cardiomyocyte cross-sectional area **(Fig. 8E)**. **Fig. 8F** demonstrates that cardiomyocyte cross-sectional area was markedly increased in HFpEF pigs vs. lean controls (median 572 µm^2^ [interquartile range, 440-743 µm^2^] vs. 385 µm^2^ [290-483 µm^2^]), indicating robust LV hypertrophy. Analysis of the distribution of individual myocytes showed that not only was the median area shifted to the right in these animals, but the spread of areas above the median was augmented, including high values that did not occur in lean controls **(Fig. 8G)**. Analysis of liver samples demonstrated significant interlobular fibrosis **(Figs. 8C and D)**.

**Fig. 8.**
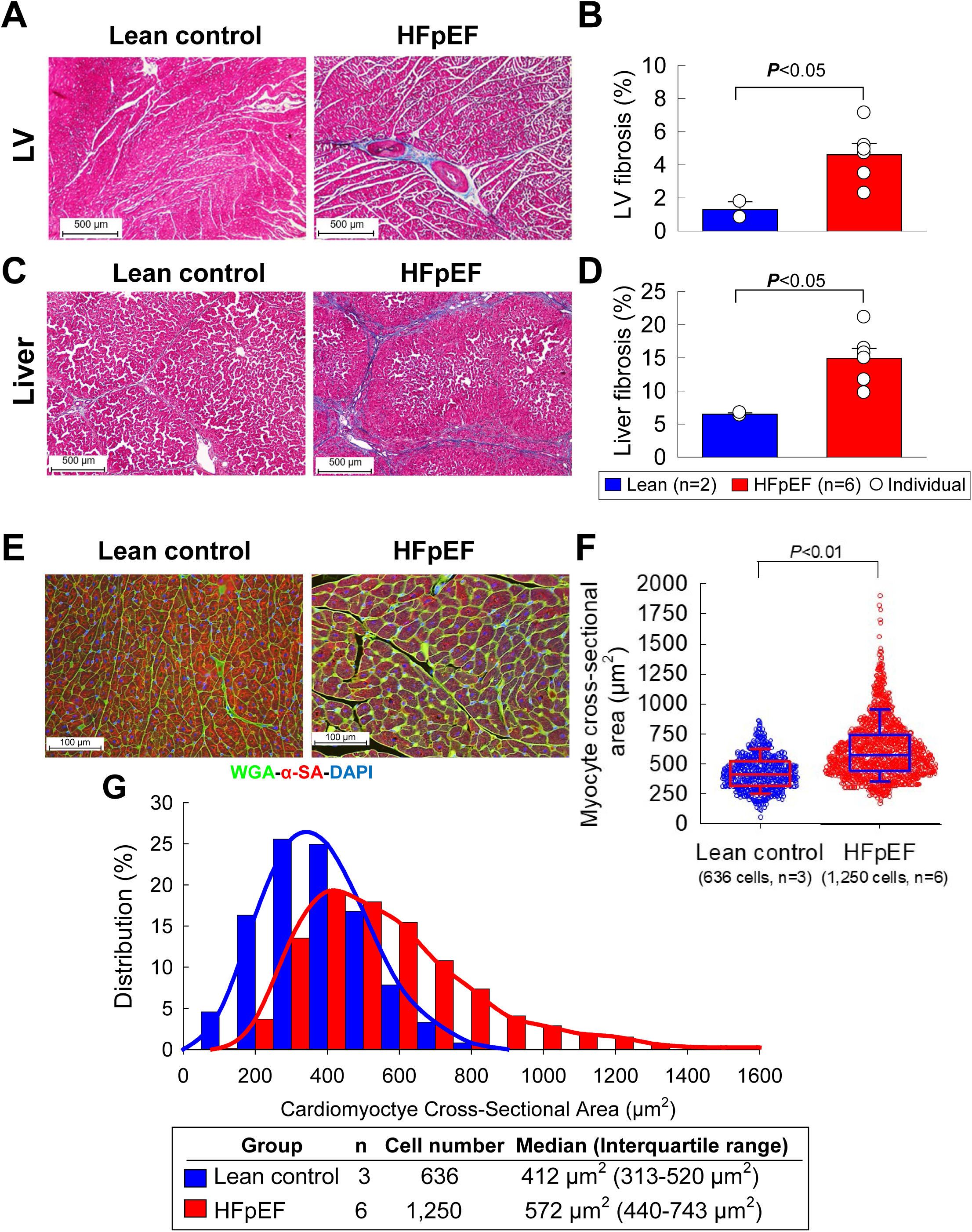
Pathologic analysis of myocardium and liver in HFpEF and lean controls pigs after 11 months of Western diet or regular diet. **A.** Representative images of myocardium stained with Masson’s trichrome in a lean and HFpEF pig. **B**. Myocardial collagen content, expressed as a percentage of total measured area. **C.** Representative images of liver stained with Masson’s trichrome in a lean and HFpEF pig. **D**. Liver collagen content, expressed as a percentage of total measured area. Data are mean ± SEM; open circles represent individual pigs. **E**. Representative images of LV myocardium stained with WGA, α-sarcomeric actin (α-SA), and DAPI from a lean control and a HFpEF pig. **F.** Cardiomyocyte cross-sectional area in the 2 cohorts. Sinaplots show individual cell distribution, boxes show the median with interquartile range (25^th^ and 75^th^ percentile), and whiskers show the 1^st^ and last decile (10^th^ and 90^th^ percentile). **G.** Distribution frequency of cardiomyocyte cross-sectional areas. Cell number is the total number of cells measured from the 2 lean controls and 6 HFpEF pigs. *P* values are shown for cross-sectional area (Wilcoxon rank-sum test) and for collagen content (unpaired Student’s t test).

## DISCUSSION

One reason for the paucity of effective therapies for HFpEF is the lack of robust and translationally relevant animal models, which represents a major obstacle both to understanding the pathophysiology of HFpEF and to developing effective treatments [7, 10, 22, 50, 52, 67, 68]. Here we describe a new, translationally relevant model of HFpEF in Ossabaw minipigs. We chose to mimic HFpEF driven by the MetS because this is the most prevalent subtype in the spectrum of human HFpEF (60-70% of all patients with HFpEF have obesity and hyperglycemia/insulin resistance) [7, 10, 22, 45, 50].

### Salient findings

After 6 months of Western diet + DOCA, Ossabaw swine exhibited marked increases in body weight, arterial blood pressure, serum cholesterol and triglycerides, and plasma glucose and insulin levels following a glucose load, indicating the development of a full MetS. These derangements were accompanied by liver fibrosis with abnormal liver function tests, suggesting the presence of MASLD [8], and reduced serum amylase levels, suggesting chronic pancreatitis [69], both of which are known to be associated with the MetS [45, 66]. Echocardiography demonstrated no change in LVEF but progressive concentric LV hypertrophy, associated with progressive left atrial dilatation. Doppler echocardiography showed that both early (E) and late atrial (A) transmitral inflow velocities increased, so that the E/A ratio did not change, which is consistent with grade 2 diastolic dysfunction in humans (“pseudonormal” LV relaxation and moderately elevated LV filling pressures) [37]. Tissue Doppler echocardiography revealed an increase in medial and lateral E/e’ ratios; the average E/e’ ratio was elevated at 2 months and continued to increase thereafter. Right heart catheterization showed an increase in central venous pressure, pulmonary arterial systolic pressure, and pulmonary capillary wedge pressure at 6 months compared with baseline. Clinically, pigs fed a Western diet for 6 months and treated with DOCA exhibited impaired exercise capacity, assessed by treadmill tests, associated with chronotropic incompetence, which is typical of the HFpEF syndrome [15, 21, 34, 43, 44, 72]. Pathologic examination of the heart showed significant myocardial fibrosis and hypertrophy.

Taken together, our observations demonstrate that this Ossabaw porcine model recapitulates the entire constellation of multiorgan comorbidities and hemodynamic, clinical, and metabolic features of human HFpEF driven by MetS: obesity, arterial hypertension, hyperlipidemia, glucose intolerance, insulin resistance, liver and pancreatic abnormalities, pulmonary hypertension, increased LV filling pressures, concentric LV hypertrophy, LV diastolic dysfunction with preserved systolic function, and impaired exercise capacity. To our knowledge, this is the first large animal study that demonstrated impaired exercise capacity in MetS-driven HFpEF, which is the hallmark of this syndrome in patients [7, 10, 22, 50–52, 68, 71]. This is also the first porcine model of HFpEF that exhibits a full MetS (documented by full metabolic characterization) including liver fibrosis and dysfunction. Because of its high translational relevance, the model described herein is well-suited for exploring the pathophysiology of MetS-driven HFpEF and the efficacy of new therapies.

### Methodological considerations

Human MetS-induced HFpEF is associated with obesity, diabetes, and hypertension [10, 45, 50]. These extracardiac comorbidities are integral in disease progression and actively contribute to HFpEF [10, 45, 50]. Thus, our pig model includes obesity, glucose intolerance, and hypertension. We used Ossabaw miniswine because these animals are genetically predisposed to MetS, recapitulate this syndrome, and thus are an ideal strain to investigate MetS-driven HFpEF and associated comorbidities [10, 12, 55, 68]. A recent review [10] concluded that “the Ossabaw swine represents a highly translatable model that develops each of the core parameters of the MetS with many of the associated comorbidities,” and that “the Ossabaw minipig stands out as the most characterized porcine model of MetS, with high relevance to human pathology.” Our results are consistent with these concepts: we found that Ossabaw pigs fed a Western diet and treated with DOCA exhibited the entire spectrum of systemic, multiorgan abnormalities associated with human MetS - central obesity, insulin resistance, glucose intolerance, dyslipidemia, hypertension, liver fibrosis, and liver and pancreatic dysfunction.

Administration of DOCA is not necessary to induce arterial hypertension in this model, since Ossabaw miniswine fed a Western diet for 6-12 months can exhibit blood pressures of ∼160/100 [13, 55]. We decided to supplement the Western diet with 1 or 2 doses of DOCA to elicit more robust hypertension and ensure that the HFpEF phenotype would develop rapidly (within a few months). This raises an intriguing point regarding the role of mineralocorticoids in the natural development of HFpEF. Unlike other pig breeds that require DOCA as a mineralocorticoid supplement [49, 52], Ossabaws may not need DOCA, because in a previous study in another Ossabaw cohort fed a similar Western diet we found the serum levels of the endogenous mineralocorticoid aldosterone to be increased six-fold (11.5 ± 1.9 compared to 2.0 ± 0.8 pg/mL in lean controls, *P*< 0.05; n=9 pigs/group). Greater serum aldosterone in MetS compared to lean Ossabaws is very likely to explain the natural increase in blood pressure, because treatment of MetS Ossabaws with the mineralocorticoid receptor blocker spironolactone decreased blood pressure to levels not significantly different from lean controls [29].

As mentioned above, liver fibrosis, abnormal liver function tests, and low serum amylase **(Table 1)** suggest liver dysfunction (MASLD) and chronic pancreatitis. These two organ derangements are an important aspect of MetS in humans [45, 66]. For example, 45-70% of patients with MetS exhibit liver dysfunction [8]. The presence of significant liver fibrosis and dysfunction sets apart our Ossabaw pig model from previous preclinical models of MetS-induced HFpEF. Indeed, MASLD has been a consistent finding in Ossabaw miniature swine, including further progression to non-alcoholic steatohepatitis [3, 27, 30] and occurrence of pancreatic steatosis with oxidative stress [18].

The two hallmarks of HFpEF in humans are i) increased LV filling pressures. and ii) reduced exercise capacity [7, 10, 22, 45, 50–52, 71]. Therefore, we assessed not only LV filling pressures (echocardiographic and hemodynamic studies), but also exercise capacity, which is the clinical correlate of LV diastolic dysfunction and represents the key clinical manifestation of HFpEF [7, 10, 22, 45, 50–52, 71]. Exercise capacity has never been measured before in swine models of MetS-induced HFpEF [28, 50, 55, 67, 68]. To assess cardiac filling pressures and diastolic function, we used two different methods: hemodynamic studies and echocardiography. These 2 independent methods yielded consistent results.

One key aspect of HFpEF is coronary vascular dysfunction, including microvascular rarefaction and decreased coronary reserve [23]. Whether these abnormalities play a causative role in the development of the HFpEF syndrome, however, is unclear [23]. While myocardial capillary density and coronary reserve were not measured in this study, previous reports have demonstrated that they are both impaired in Ossabaw swine with MetS [33, 65].

Ossabaw miniswine have a unique genetic background that not only predisposes them to obesity, MetS, and atherosclerosis [10, 12, 55, 68], but also results in another characteristic that may promote development of HFpEF, i.e., a predisposition to a vascular reactivity profile that would result in lower coronary blood flow compared with other pigs strains such as the Gӧttingen miniature pig. This vascular reactivity profile was observed in *ex vivo* mesenteric arteries: vessels isolated from Ossabaw pigs exhibited greater vasoconstriction to norepinephrine, less endothelium-dependent vasodilation to carbachol, and less endothelium-independent vasodilation to nitroprusside than those from Gӧttingen pigs [14]. The unique genetic background of Ossabaw miniswine also manifests itself in an inability to develop a cardioprotective phenotype in response to ischemic preconditioning [25, 31, 32]. The full sequencing, assembly, and annotation of the Ossabaw miniature pig genome provides an outstanding opportunity to determine specific genes underlying the predisposition to abnormal vasomotor function, non-responsiveness to cardioprotection by ischemic preconditioning, and HFpEF [5, 16, 70].

### Exercise tolerance

Decreased exercise tolerance in pigs is generally defined by any of the following characteristics: 1) excessive HR, 2) abnormal gait, 3) labored breathing, 4) fall on the treadmill, 5) excessive vocalizing, and 6) decreased exercise HR [44]. In this study, decreased exercise tolerance was evident in 7 of the 9 HFpEF pigs, requiring completely stopping and/or decreasing workload **(Fig. 7)**. In contrast to lean pigs that all completed the 10-min test, only 2 of 9 HFpEF pigs were able to complete the 10-min fast walking phase of the protocol, the mean duration being 6.3 ± 0.9 min (*P*<0.002 vs. baseline; **Fig. 7**).

The HR response was also abnormal in HFpEF pigs. it should be noted that the 10-min treadmill test at 7.7 km/h is only a fast walk; thus, the average HR of 117/min at baseline was only ∼47% of the predicted maximum HR of ∼250/min. Obesity and metabolic syndrome of this severity should cause a statistically significant, but moderate, increase in HR to compensate for the increased mass of the pigs. This is a fundamental premise of exercise physiology [44]. Two of the 9 HFpEF pigs had dangerously high HR responses to the test, which prompted us to decrease the workload. The other 7 HFpEF pigs showed a trend toward decreased HR responses to the test compared to their healthy baseline and to 3 lean controls. Taken together, the results of the treadmill exercise test strongly support a diagnosis of decreased exercise tolerance in all HFpEF pigs and chronotropic incompetence in the majority of them.

### Previous studies of HFpEF

Previous studies of HFpEF in rodents have used mice treated with NO synthase inhibitors to induce hypertension [17] and spontaneously hypertensive rats [11, 19]. Rodent models are useful to interrogate the molecular mechanisms underlying disease; however, for clinical translation to be possible, rodent data need to be tested in large animals before embarking in clinical trials with their attendant costs, risks, and efforts.

**Table 2** summarizes the previous porcine models of HFpEF [9, 20, 36, 41, 46, 47, 49, 52, 53, 67]. Most of these studies have used animals made hypertensive with angiotensin II [20], DOCA [46], or partial nephrectomy [47] or subjected to aortic constriction [9, 41]. The major differences between these previous models and the present study are the absence of either MetS or full metabolic characterization, the absence of hepatic fibrosis and dysfunction (present in 45-70% of patients with HFpEF [45, 69]), and the fact that exercise tolerance was not measured [9, 20, 36, 41, 46, 47, 49, 52, 53, 67]. The pig strain is also important. Many previous studies have used Landrace, Yorkshire, and other full-sized pigs [9, 20, 36, 46, 47, 49, 53]; these strains are problematic for long-term testing of therapies because of the rapid weight gain of the animals over time.

**Table 2.**
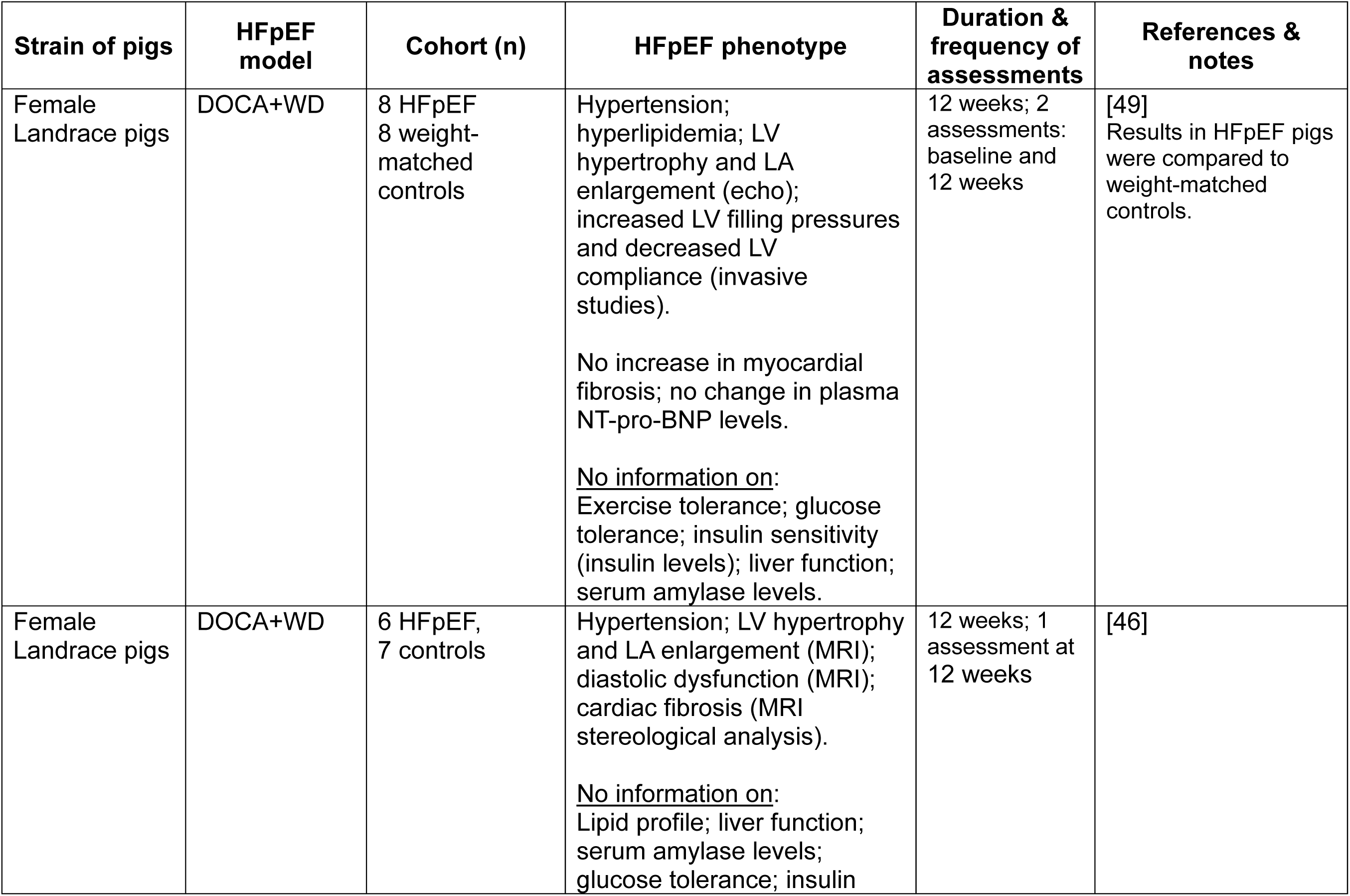
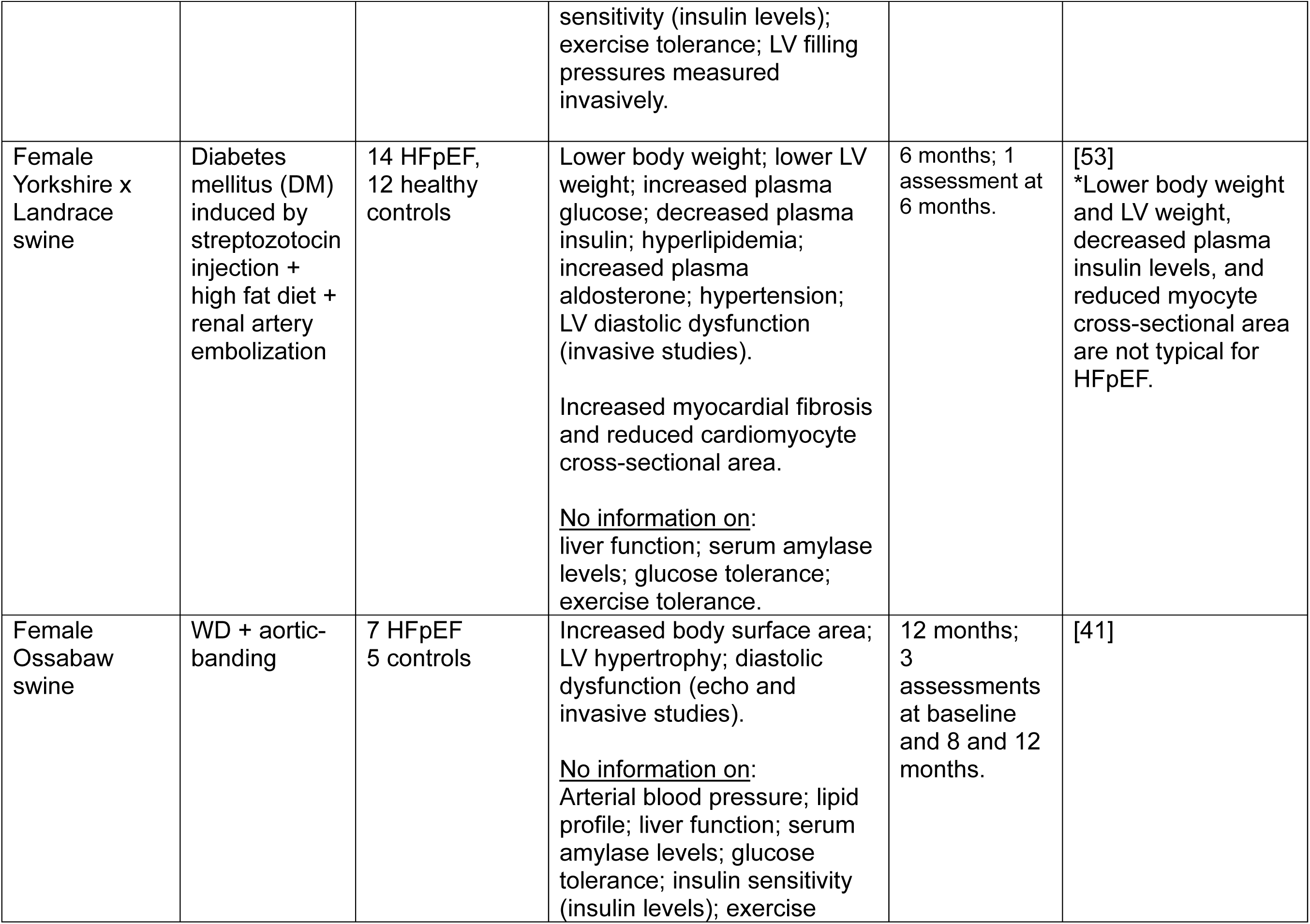
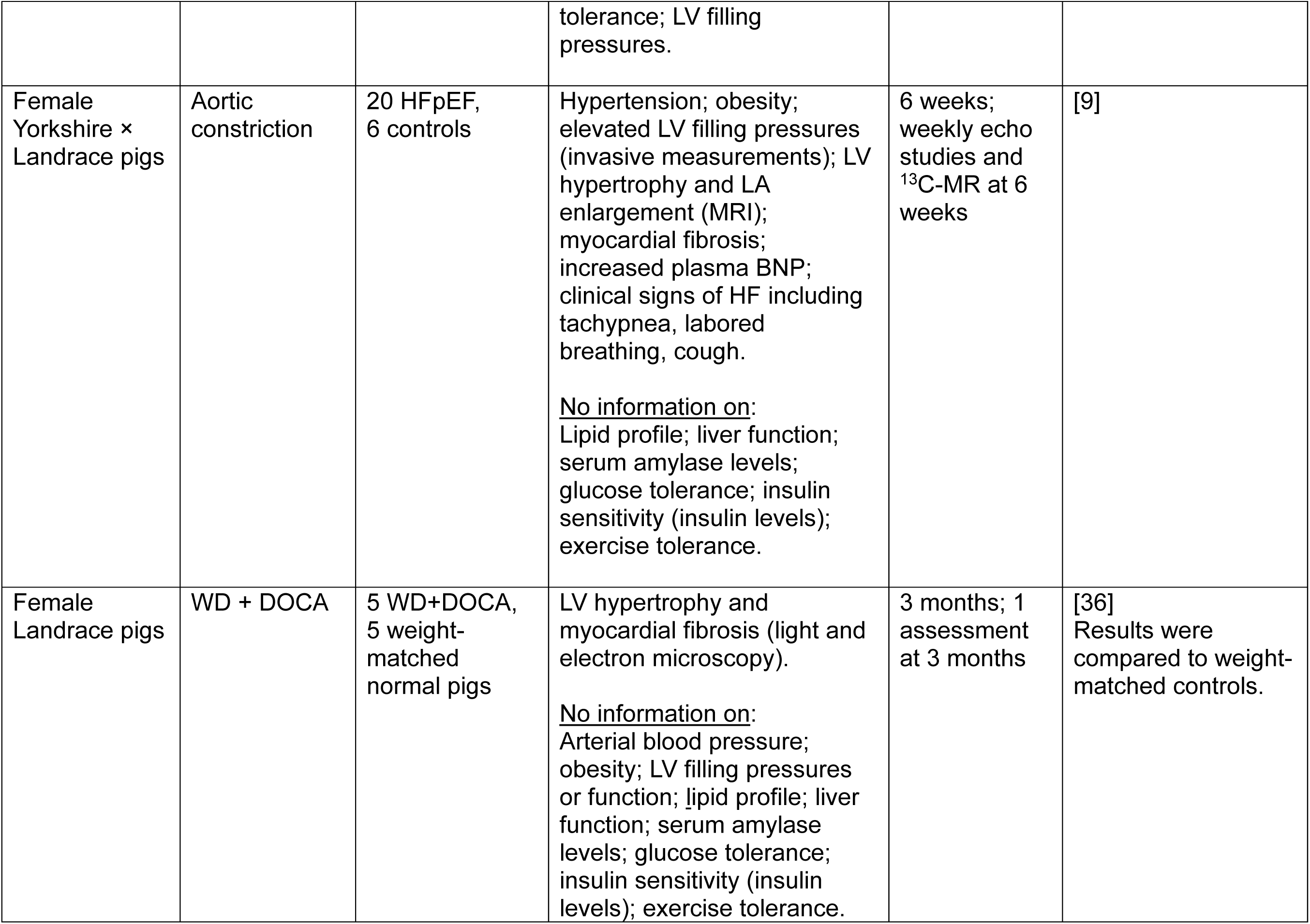
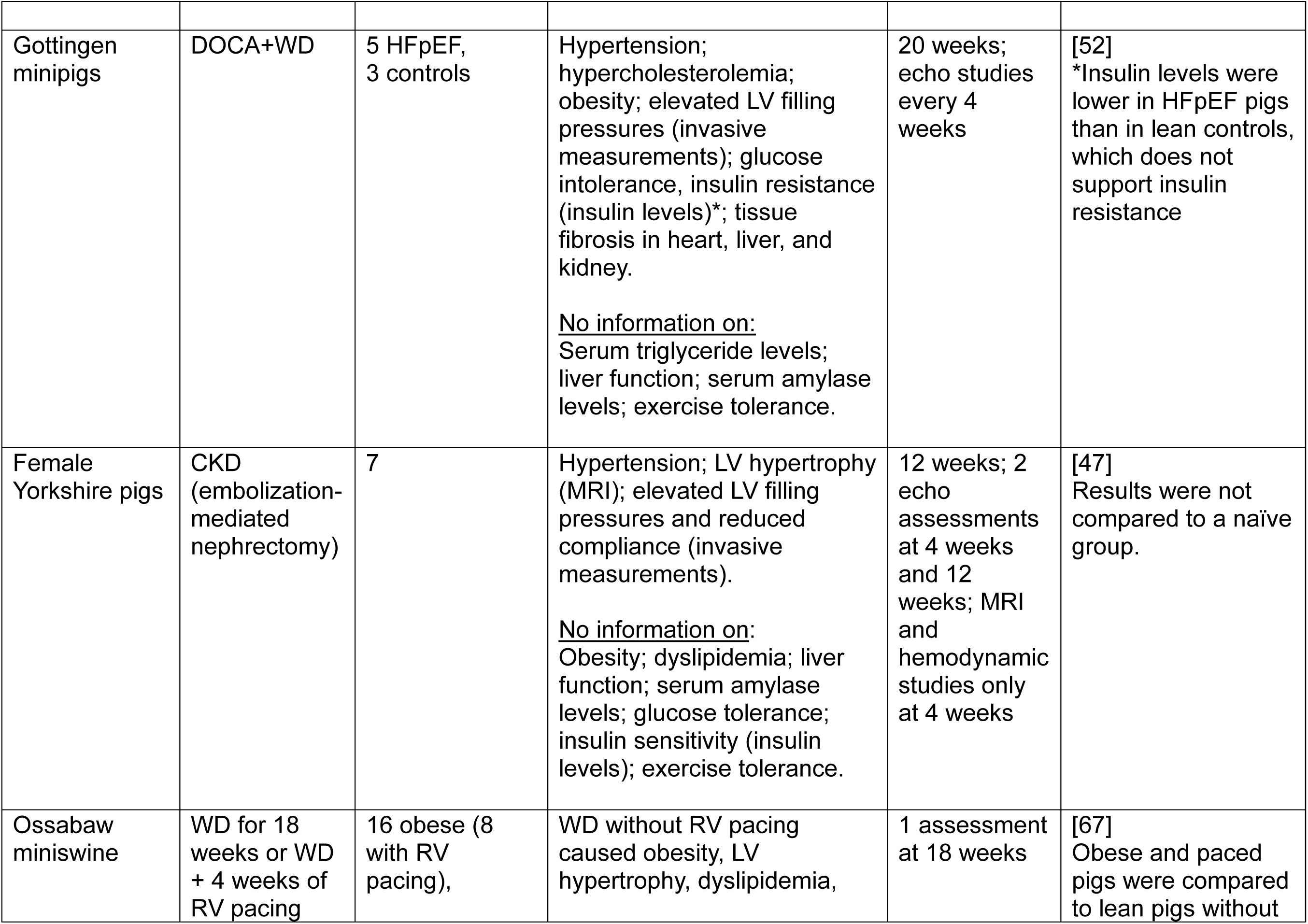
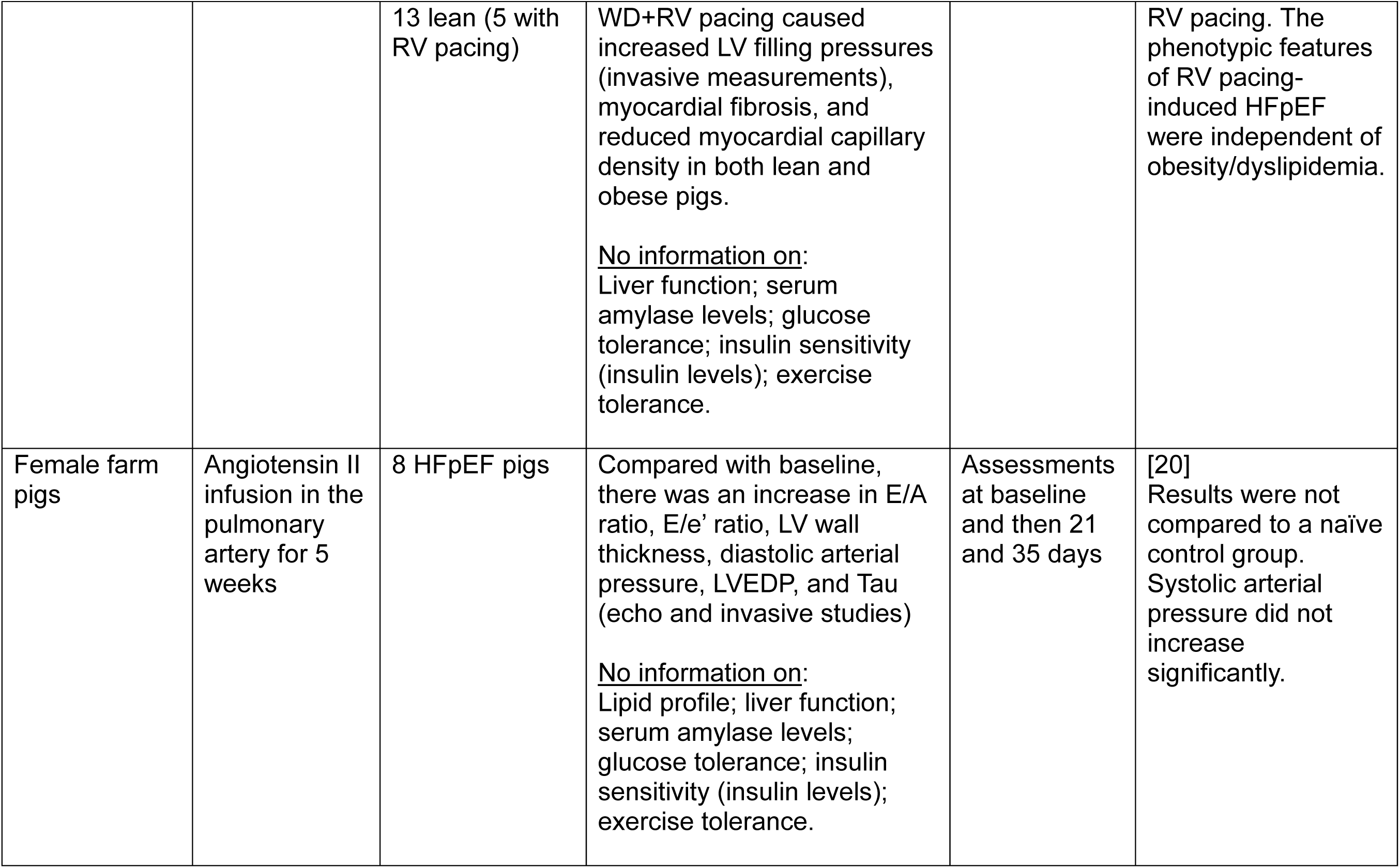
Summary of previous HFpEF models in pigs.

| **Strain of pigs** | **HFpEF model** | **Cohort (n)** | **HFpEF phenotype** | **Duration & frequency of assessments** | **References & notes** |
| --- | --- | --- | --- | --- | --- |
| Female Landrace pigs | DOCA+WD | 8 HFpEF<br>8 weight-matched controls | Hypertension; hyperlipidemia; LV hypertrophy and LA enlargement (echo); increased LV filling pressures and decreased LV compliance (invasive studies).<br>No increase in myocardial fibrosis; no change in plasma NT-pro-BNP levels.<br><u>No information on:</u><br>Exercise tolerance; glucose tolerance; insulin sensitivity (insulin levels); liver function; serum amylase levels. | 12 weeks; 2 assessments: baseline and 12 weeks | [49]<br>Results in HFpEF pigs were compared to weight-matched controls. |
| Female Landrace pigs | DOCA+WD | 6 HFpEF,<br>7 controls | Hypertension; LV hypertrophy and LA enlargement (MRI); diastolic dysfunction (MRI); cardiac fibrosis (MRI stereological analysis).<br><u>No information on:</u><br>Lipid profile; liver function; serum amylase levels; glucose tolerance; insulin<br>sensitivity (insulin levels); exercise tolerance; LV filling pressures measured invasively. | 12 weeks; 1 assessment at 12 weeks | [46] |
| Female Yorkshire x Landrace swine | Diabetes mellitus (DM) induced by streptozotocin injection + high fat diet + renal artery embolization | 14 HFpEF, 12 healthy controls | Lower body weight; lower LV weight; increased plasma glucose; decreased plasma insulin; hyperlipidemia; increased plasma aldosterone; hypertension; LV diastolic dysfunction (invasive studies).<br>Increased myocardial fibrosis and reduced cardiomyocyte cross-sectional area.<br><u>No information on:</u> liver function; serum amylase levels; glucose tolerance; exercise tolerance. | 6 months; 1 assessment at 6 months. | [53]<br>*Lower body weight and LV weight, decreased plasma insulin levels, and reduced myocyte cross-sectional area are not typical for HFpEF. |
| Female Ossabaw swine | WD + aortic-banding | 7 HFpEF 5 controls | Increased body surface area; LV hypertrophy; diastolic dysfunction (echo and invasive studies).<br><u>No information on:</u> Arterial blood pressure; lipid profile; liver function; serum amylase levels; glucose tolerance; insulin sensitivity (insulin levels); exercise<br>tolerance; LV filling pressures. | 12 months; 3 assessments at baseline and 8 and 12 months. | [41] |
| Female Yorkshire × Landrace pigs | Aortic constriction | 20 HFpEF, 6 controls | Hypertension; obesity; elevated LV filling pressures (invasive measurements); LV hypertrophy and LA enlargement (MRI); myocardial fibrosis; increased plasma BNP; clinical signs of HF including tachypnea, labored breathing, cough.<br><u>No information on:</u><br>Lipid profile; liver function; serum amylase levels; glucose tolerance; insulin sensitivity (insulin levels); exercise tolerance. | 6 weeks; weekly echo studies and <sup>13</sup>C-MR at 6 weeks | [9] |
| Female Landrace pigs | WD + DOCA | 5 WD+DOCA, 5 weight-matched normal pigs | LV hypertrophy and myocardial fibrosis (light and electron microscopy).<br><u>No information on:</u><br>Arterial blood pressure; obesity; LV filling pressures or function; lipid profile; liver function; serum amylase levels; glucose tolerance; insulin sensitivity (insulin levels); exercise tolerance. | 3 months; 1 assessment at 3 months | [36]<br>Results were compared to weight-matched controls. |
| Gottingen minipigs | DOCA+WD | 5 HFpEF, 3 controls | Hypertension; hypercholesterolemia; obesity; elevated LV filling pressures (invasive measurements); glucose intolerance, insulin resistance (insulin levels)*; tissue fibrosis in heart, liver, and kidney.<br><u>No information on:</u><br>Serum triglyceride levels; liver function; serum amylase levels; exercise tolerance. | 20 weeks; echo studies every 4 weeks | [52]<br>*Insulin levels were lower in HFpEF pigs than in lean controls, which does not support insulin resistance |
| Female Yorkshire pigs | CKD (embolization-mediated nephrectomy) | 7 | Hypertension; LV hypertrophy (MRI); elevated LV filling pressures and reduced compliance (invasive measurements).<br><u>No information on:</u><br>Obesity; dyslipidemia; liver function; serum amylase levels; glucose tolerance; insulin sensitivity (insulin levels); exercise tolerance. | 12 weeks; 2 echo assessments at 4 weeks and 12 weeks; MRI and hemodynamic studies only at 4 weeks | [47]<br>Results were not compared to a naïve group. |
| Ossabaw miniswine | WD for 18 weeks or WD + 4 weeks of RV pacing | 16 obese (8 with RV pacing), | WD without RV pacing caused obesity, LV hypertrophy, dyslipidemia, | 1 assessment at 18 weeks | [67]<br>Obese and paced pigs were compared to lean pigs without |
|  |  | 13 lean (5 with RV pacing) | WD+RV pacing caused increased LV filling pressures (invasive measurements), myocardial fibrosis, and reduced myocardial capillary density in both lean and obese pigs.<br><u>No information on:</u><br>Liver function; serum amylase levels; glucose tolerance; insulin sensitivity (insulin levels); exercise tolerance. |  | RV pacing. The phenotypic features of RV pacing-induced HFpEF were independent of obesity/dyslipidemia. |
| Female farm pigs | Angiotensin II infusion in the pulmonary artery for 5 weeks | 8 HFpEF pigs | Compared with baseline, there was an increase in E/A ratio, E/e' ratio, LV wall thickness, diastolic arterial pressure, LVEDP, and Tau (echo and invasive studies)<br><u>No information on:</u><br>Lipid profile; liver function; serum amylase levels; glucose tolerance; insulin sensitivity (insulin levels); exercise tolerance. | Assessments at baseline and then 21 and 35 days | [20]<br>Results were not compared to a naïve control group. Systolic arterial pressure did not increase significantly. |

HFpEF has been described in Ossabaw swine after chronic atrial pacing [67]. Major differences from our model are clinical relevance (tachycardia is rarely, if ever, a clinical cause of HFpEF), absence of exercise testing, and the fact that minimal metabolic characterization was carried out and that blood pressure and glucose were measured under general anesthesia. Gottingen miniswine have also been reported to develop HFpEF when fed a Western diet with DOCA [52], but unlike Ossabaw miniswine, they do not exhibit liver fibrosis/dysfunction [12, 50] and exercise tolerance was not assessed [52]; furthermore, there are no reports on coronary microvascular dysfunction in this model [55], and hypertension and coronary macrovascular dysfunction have only been noted in one report [52]. A porcine model of HFpEF has been produced by a 5/6 nephrectomy [47]. Major differences vs. the present model are, again, the absence of MetS and the fact that clinical endpoints (e.g., exercise tolerance) were not evaluated [47].

### Study limitations

The small number of animals in the lean control group (n=2-3) reflects the necessity to limit the costs of these complex and long-lasting studies, which required longitudinal follow-up for many months and a battery of hemodynamic, echocardiographic, functional, metabolic, and biochemical assessments. The small number of lean controls also reflects the fact that the Ossabaw miniswine has been well characterized [10, 12, 55, 68]; the results in controls were sufficiently consistent that the differences described herein reached statistical significance despite the small sample size. LV filling pressures were not measured directly with an LV catheter; however, we measured PCWP, which reflects LV filling pressures closely. Histologic data **(Fig. 8)** were obtained at 11 months rather than at 6 months. This was necessitated by the fact that pigs were sacrificed at 11 months after starting the Western diet (the last 5 months were used for pilot studies of experimental therapies). However, the histologic data presented here were obtained only in control animals that did not receive any therapy, and thus are relevant to the phenotypic features of this HFpEF model.

### Conclusions

We describe a porcine model that recapitulates the multiorgan involvement and the constellation of comorbidities and hemodynamic, clinical, and metabolic features of human HFpEF driven by MetS, namely, obesity, arterial hypertension, hyperlipidemia, glucose intolerance, insulin resistance, liver and pancreatic dysfunction, pulmonary hypertension, increased LV filling pressures, concentric LV hypertrophy, LV diastolic dysfunction with preserved systolic function, and impaired exercise tolerance. Importantly, we assessed the clinical correlate of diastolic dysfunction, i.e., exercise capacity measured with treadmill tests, which has never been measured before in swine models of MetS-induced HFpEF [28, 49, 50, 52, 55, 68]. The model described herein is unique in that it is the only porcine model published heretofore that exhibits all of the following components: a full MetS, a full metabolic characterization, liver fibrosis and dysfunction, and reduced exercise tolerance. Although this model is expensive, complex, and time-intensive, it has high translational relevance and should be useful for preclinical testing of new therapies for HFpEF, including devices.

## Supporting information

Model of HFpEF

## ACKNOWLEDMENTS

This study was supported by Department of Defense grant HT94252310109 (R.B.).

## Conflict of Interest Disclosures

The authors declare no conflict of interest.

## Notes

### Competing Interest Statement

The authors have declared no competing interest.

