## Supplementary material for "A New Model of Heart Failure with Preserved Ejection Fraction Induced by Metabolic Syndrome in Ossabaw Miniature Swine": Model of HFpEF

**Supplementary Table 1. Echocardiographic variables**

| | Time | Lean (n=3) | | HFpEF (n=10) | |
|---|---|---|---|---|---|
| | | Mean | SEM | Mean | SEM |
| **LVEDV** (ml) | Baseline | 69.33 | 7.63 | 63.87 | 5.56 |
| | 2-month | 64.26 | 0.49 | 72.91 | 8.53 |
| | 4-month | 69.00 | 1.55 | 78.20 | 6.59 |
| | 6-month | 72.59 | 9.03 | 77.84 | 6.15 |
| **LVESV** (ml) | Baseline | 26.68 | 6.12 | 26.49 | 3.08 |
| | 2-month | 26.55 | 2.58 | 29.55 | 3.81 |
| | 4-month | 33.75 | 2.40 | 31.57 | 2.89 |
| | 6-month | 32.81 | 4.20 | 32.69 | 4.27 |
| **SV** (ml) | Baseline | 42.65 | 2.02 | 37.32 | 2.96 |
| | 2-month | 37.70 | 2.80 | 41.29 | 4.76 |
| | 4-month | 35.25 | 0.90 | 46.63 | 4.32 |
| | 6-month | 39.78 | 5.33 | 45.15 | 2.59 |
| **EF** (%) | Baseline | 62.53 | 4.77 | 59.22 | 2.21 |
| | 2-month | 58.65 | 4.14 | 59.00 | 1.82 |
| | 4-month | 51.20 | 2.40 | 59.50 | 2.08 |
| | 6-month | 54.68 | 2.11 | 59.17 | 2.39 |
| **IVSd** (cm) | Baseline | 0.93 | 0.04 | 0.97 | 0.03 |
| | 2-month | 0.97 | 0.04 | 1.17 | 0.06 |
| | 4-month | 0.92 | 0.01 | 1.28 | 0.05 |
| | 6-month | 0.87 | 0.04 | 1.27 | 0.05 |
| **LVIDd** (cm) | Baseline | 4.01 | 0.17 | 3.99 | 0.14 |
| | 2-month | 3.87 | 0.07 | 3.70 | 0.16 |
| | 4-month | 4.07 | 0.23 | 3.83 | 0.18 |
| | 6-month | 4.23 | 0.14 | 3.64 | 0.14 |
| **PWd** (cm) | Baseline | 0.92 | 0.06 | 0.88 | 0.05 |
| | 2-month | 0.84 | 0.09 | 1.11 | 0.07 |
| | 4-month | 0.89 | 0.01 | 1.22 | 0.05 |
| | 6-month | 0.87 | 0.05 | 1.26 | 0.06 |
| **IVSs** (cm) | Baseline | 1.30 | 0.08 | 1.42 | 0.09 |
| | 2-month | 1.20 | 0.08 | 1.53 | 0.13 |
| | 4-month | 1.18 | 0.03 | 1.66 | 0.09 |
| | 6-month | 1.19 | 0.02 | 1.61 | 0.06 |
| **LVIDs** (cm) | Baseline | 2.64 | 0.27 | 2.53 | 0.18 |
| | 2-month | 2.71 | 0.12 | 2.21 | 0.23 |
| | 4-month | 2.90 | 0.36 | 2.40 | 0.17 |
| | 6-month | 2.88 | 0.15 | 2.30 | 0.21 |
| **PWs** (cm) | Baseline | 1.26 | 0.17 | 1.29 | 0.10 |
| | 2-month | 1.19 | 0.10 | 1.54 | 0.11 |

| | | | | | |
|---|---|---|---|---|---|
| | 4-month | 1.18 | 0.04 | 1.70 | 0.09 |
| | 6-month | 1.17 | 0.06 | 1.63 | 0.07 |
| **LV Mass** (g) | Baseline | 114.82 | 7.29 | 112.95 | 5.71 |
| | 2-month | 104.75 | 6.67 | 135.03 | 8.63 |
| | 4-month | 111.01 | 8.05 | 168.18 | 11.94 |
| | 6-month | 124.46 | 2.13 | 165.25 | 11.82 |
| **Teicholz EDV** (ml) | Baseline | 70.44 | 7.62 | 70.52 | 5.55 |
| | 2-month | 64.81 | 2.76 | 59.52 | 6.30 |
| | 4-month | 73.71 | 9.63 | 65.44 | 7.65 |
| | 6-month | 80.11 | 6.53 | 56.54 | 5.21 |
| **Teicholz** ESV (ml) | Baseline | 26.57 | 6.80 | 24.79 | 3.69 |
| | 2-month | 27.46 | 3.01 | 19.25 | 4.22 |
| | 4-month | 33.97 | 10.33 | 22.04 | 3.94 |
| | 6-month | 32.05 | 4.03 | 20.43 | 4.42 |
| **Teicholz SV** (ml) | Baseline | 43.87 | 2.00 | 45.73 | 4.61 |
| | 2-month | 37.35 | 4.32 | 40.18 | 4.28 |
| | 4-month | 39.74 | 5.60 | 43.41 | 5.66 |
| | 6-month | 48.06 | 5.33 | 36.20 | 3.27 |
| **Teicholz EF (**%) | Baseline | 63.34 | 5.60 | 64.87 | 4.55 |
| | 2-month | 57.20 | 5.00 | 69.22 | 6.01 |
| | 4-month | 55.56 | 8.93 | 66.35 | 4.29 |
| | 6-month | 59.88 | 4.33 | 66.70 | 5.13 |
| **FS** (%) | Baseline | 34.22 | 4.04 | 36.54 | 4.02 |
| | 2-month | 29.91 | 3.40 | 40.84 | 5.05 |
| | 4-month | 29.13 | 5.54 | 37.10 | 3.36 |
| | 6-month | 31.76 | 3.10 | 37.43 | 3.64 |
| **RWT** (%) | Baseline | 0.46 | 0.03 | 0.45 | 0.04 |
| | 2-month | 0.43 | 0.06 | 0.62 | 0.05 |
| | 4-month | 0.44 | 0.03 | 0.66 | 0.05 |
| | 6-month | 0.41 | 0.04 | 0.70 | 0.05 |
| **E** (cm/s) | Baseline | 61.99 | 3.11 | 50.32 | 1.60 |
| | 2-month | 58.22 | 1.96 | 57.99 | 3.08 |
| | 4-month | 52.86 | 3.74 | 63.86 | 4.83 |
| | 6-month | 46.78 | 1.13 | 60.32 | 4.50 |
| **A** (cm/s) | Baseline | 35.91 | 7.38 | 32.49 | 1.25 |
| | 2-month | 30.03 | 5.34 | 36.36 | 2.38 |
| | 4-month | 24.10 | 3.88 | 40.94 | 4.52 |
| | 6-month | 24.68 | 2.84 | 38.99 | 3.68 |

| | | | | | |
|---|---|---|---|---|---|
| **E/A** | Baseline | 1.86 | 0.33 | 1.57 | 0.08 |
| | 2-month | 2.03 | 0.26 | 1.62 | 0.06 |
| | 4-month | 2.27 | 0.23 | 1.67 | 0.17 |
| | 6-month | 1.96 | 0.27 | 1.61 | 0.13 |
| **IVRT (**s) | Baseline | 0.07 | 0.00 | 0.07 | 0.00 |
| | 2-month | 0.08 | 0.00 | 0.07 | 0.01 |
| | 4-month | 0.08 | 0.00 | 0.06 | 0.00 |
| | 6-month | 0.09 | 0.01 | 0.07 | 0.01 |
| **R-R interval** (s) | Baseline | 0.83 | 0.07 | 0.76 | 0.06 |
| | 2-month | 0.88 | 0.14 | 0.72 | 0.05 |
| | 4-month | 0.90 | 0.06 | 0.69 | 0.05 |
| | 6-month | 1.29 | 0.05 | 0.73 | 0.07 |
| **HR** (bpm) | Baseline | 73.68 | 6.95 | 82.53 | 6.16 |
| | 2-month | 72.19 | 13.29 | 88.49 | 6.71 |
| | 4-month | 67.64 | 5.14 | 92.11 | 7.62 |
| | 6-month | 46.80 | 2.03 | 91.38 | 11.86 |
| **e' SA (**cm/s) | Baseline | -9.28 | 1.53 | -9.62 | 0.71 |
| | 2-month | -8.55 | 1.04 | -7.87 | 0.34 |
| | 4-month | -6.88 | 0.98 | -7.19 | 0.53 |
| | 6-month | -6.67 | 0.47 | -6.73 | 0.51 |
| **a' SA** (cm/s) | Baseline | -4.44 | 0.90 | -5.84 | 0.49 |
| | 2-month | -5.88 | 0.79 | -5.48 | 0.35 |
| | 4-month | -4.02 | 0.13 | -5.13 | 0.37 |
| | 6-month | -4.30 | 0.92 | -4.87 | 0.54 |
| **SA (E/e')** | Baseline | 6.94 | 0.79 | 5.47 | 0.43 |
| | 2-month | 7.01 | 0.85 | 7.40 | 0.34 |
| | 4-month | 7.87 | 0.66 | 8.95 | 0.40 |
| | 6-month | 7.07 | 0.38 | 9.25 | 0.76 |
| **e' LA (**cm/s) | Baseline | -10.19 | 2.03 | -9.11 | 0.69 |
| | 2-month | -9.82 | 1.04 | -8.17 | 0.55 |
| | 4-month | -8.74 | 1.10 | -7.24 | 0.48 |
| | 6-month | -7.22 | 0.62 | -7.59 | 0.54 |
| **a' LA** (cm/s) | Baseline | -6.41 | 1.23 | -6.11 | 0.70 |
| | 2-month | -6.14 | 0.31 | -6.01 | 0.54 |
| | 4-month | -6.68 | 0.87 | -5.51 | 0.56 |
| | 6-month | -4.77 | 0.31 | -5.36 | 0.32 |
| **LA (E/e')** | Baseline | 6.46 | 0.97 | 5.88 | 0.58 |
| | 2-month | 6.06 | 0.67 | 7.27 | 0.48 |
| | 4-month | 6.15 | 0.40 | 9.02 | 0.74 |
| | 6-month | 6.56 | 0.47 | 8.00 | 0.42 |
| **e' (mean)** | Baseline | -9.73 | 1.28 | -9.38 | 0.65 |

| | | | | | |
|---|---|---|---|---|---|
| | 2-month | -9.19 | 1.00 | -8.02 | 0.38 |
| | 4-month | -7.81 | 1.03 | -7.21 | 0.43 |
| | 6-month | -6.94 | 0.54 | -7.16 | 0.45 |
| **a' (mean)** | Baseline | -5.43 | 1.05 | -5.98 | 0.52 |
| | 2-month | -6.01 | 0.49 | -5.75 | 0.40 |
| | 4-month | -5.35 | 0.38 | -5.32 | 0.40 |
| | 6-month | -4.54 | 0.60 | -5.12 | 0.32 |
| **E/e' (**mean) | Baseline | 6.53 | 0.62 | 5.62 | 0.47 |
| | 2-month | 6.49 | 0.73 | 7.27 | 0.32 |
| | 4-month | 6.90 | 0.50 | 8.86 | 0.45 |
| | 6-month | 6.80 | 0.43 | 8.50 | 0.52 |
| **LA AreaED** (cm2) | Baseline | 9.73 | 0.71 | 10.29 | 0.61 |
| | 2-month | 10.28 | 0.74 | 11.58 | 0.71 |
| | 4-month | 9.91 | 0.94 | 11.45 | 0.70 |
| | 6-month | 10.44 | 0.19 | 13.53 | 0.59 |
| **LAFAC** (%) | Baseline | 42.96 | 0.82 | 46.49 | 2.27 |
| | 2-month | 46.38 | 2.22 | 30.39 | 1.63 |
| | 4-month | 39.53 | 2.23 | 28.43 | 1.73 |
| | 6-month | 42.40 | 1.67 | 25.91 | 2.19 |

[a] *P*<0.05, [b] *P*<0.01 vs. baseline within group; ***** *P*<0.05, ****** P<0.01 vs. lean control

**LVEDV**, left ventricular end-diastolic volume; **LVESV**, left ventricular end-systolic volume; **SV**, stroke volume; **EF**, ejection fraction; **IVSd**, interventricular septum thickness at end-diastole; **IVSs**, interventricular septum thickness at end-systole; **LVIDd**, left ventricular internal dimension at end-diastole; **LVIDs**, left ventricular internal dimension at end-systole; **PWd**, posterior wall thickness at end-diastole; **PWs**, posterior wall thickness at end-systole; **Teicholz EDV**, left ventricular end-diastolic volume measured by the Teicholz method; **Teicholz ESV**, left ventricular end-systolic volume measured by the Teicholz method; **Teicholz SV**, stroke volume measured by the Teicholz method; **FS**, fractional shortening; **RWT**, relative left ventricular wall thickness; **IVRT**, isovolumic relaxation time; **HR**, heart rate; **e' or a' SA**, median (septal) annulus tissue Doppler e' or a'; **e' or a' LA**, lateral annulus tissue Doppler e' or a'; **LA area$_{ED}$**, left atrial area at end-diastole; **LAFAC**, left atrial fractional area change.

**Supplementary Table 2. Right heart pressures**

| | Time | Lean (n=2) | | HFpEF (n=9) | |
|---|---|---|---|---|---|
| | | Mean | SEM | Mean | SEM |
| CVP (mmHg) | Baseline | 7.2 | 2.2 | 6.4 | 0.9 |
| | 6-month | 6.4 | 0.7 | 10.0 | 1.9[a] |
| RVSP (mmHg) | Baseline | 30.0 | 6.0 | 26.3 | 2.0 |
| | 6-month | 23.0 | 1.2 | 38.3 | 2.0 [b**] |
| PASP (mmHg) | Baseline | 29.0 | 5.0 | 24.6 | 1.8 |
| | 6-month | 22.3 | 0.9 | 35.3 | 1.7 [b**] |
| PCWP (mmHg) | Baseline | 8.5 | 3.2 | 9.4 | 0.9 |
| | 6-month | 9.0 | 1.9 | 16.6 | 1.7 [b**] |

[a] $P<0.05$, [b] $P<0.01$ vs. baseline within group; * $P<0.05$, ** $P<0.01$ vs. lean control

**CVP**, central venous pressure; **RVSP**, right ventricular systolic pressure; **PASP**, pulmonary arterial systolic pressure; **PCWP**, pulmonary capillary wedge pressure.
